# Characterizing Tissue Structures from Spatial Omics with Spatial Cellular Graph Partition

**DOI:** 10.1101/2023.09.05.556133

**Authors:** Zhenqin Wu, Ayano Kondo, Monee McGrady, Ethan A. G. Baker, Eric Wu, Maha K. Rahim, Nathan A. Bracey, Vivek Charu, Raymond J. Cho, Jeffrey B. Cheng, Maryam Afkarian, James Zou, Aaron T. Mayer, Alexandro E. Trevino

## Abstract

Spatial transcriptomic and proteomic measurements enable high-dimensional characterization of tissues. However, understanding organizations of cells at different spatial scales and extracting tissue structures of interest remain challenging tasks that require extensive human annotations. To address this need for consistent identification of tissue structures, in this work, we present a novel annotation method Spatial Cellular Graph Partitioning (SCGP) that allows unsupervised identification of tissue structures that reflect the anatomical and functional units of human tissues. We further present a reference-query extension pipeline SCGP-Extension that enables the generalization of existing reference tissue structures to previously unseen samples. Our experiments demonstrate reliable and robust partitionings of both spatial transcriptomics and proteomics datasets encompassing different tissue types and profiling techniques. Downstream analysis on SCGP-identified tissue structures reveals disease-relevant insights regarding diabetic kidney disease and skin disorder, underscoring its potential in facilitating spatial analysis and driving new discoveries.

## Introduction

All human organs exhibit characteristic tissue structures that are required for homeostasis and function. These structures are diverse in form, scale and function and are typically composed of multiple cell types organized into spatial patterns. Disruptions to these structures are usually indicative of a disease process^1,2^. Recent advances in *in situ* molecular profiling techniques, including spatial transcriptomics^3–7^ and proteomics^8–11^ techniques, have allowed us to observe detailed molecular phenotypes and cell states in tissue context, enabling the exploration of interactions between entities at different spatial scales, ranging from cells^12^ and cellular neighborhoods^13,14^ to tissue organization^15^ and patient-level characteristics^16,17^. However, connecting observations of multi-scale tissue structures to molecular pathways and cellular communities remains a challenging task.

Many existing analysis pipelines^18,19^ for spatially resolved data are cell-centric: cells are identified and annotated through segmentation^20^, clustering, and classification^21,22^. Uncovering the multi-scale tissue structures in parallel with the cell annotations can enable more comprehensive understanding of tissues and facilitates downstream analysis focused on specific structures-of-interest. Moreover, when comparing diverse samples from different experiments or with different disease conditions, it is necessary to consistently recognize the same set of tissue structures across all samples. In practice, this is best accomplished by extending annotations on a well-studied set of samples to previously unseen samples, a process we refer to as “generalization”.

In recent studies, computational methods integrating molecular profiling with spatial information have been proposed. Some of these methods aim to improve analysis of cell-level characteristics, such as better cell type prediction^23,24^ and intercellular communication modeling^12,25^. Another line of research focuses on annotating larger structures or spatial domains, exploring their interactions and disease-relevance. Such annotations are performed based on clustering of cell type composition^13,26^ or locally-smoothed cell features^15^, topic modeling^27^, Bayesian modeling^28^, optimal transport^29^, graph Fourier transform^30^, and graph neural networks^16,31,32^. Many of these methods are unsupervised and lack the ability to generalize. When new data is introduced, model retraining or re-fitting is necessary to annotate unseen data. Consequently, downstream analysis on structures-of-interest are restricted to only the training/fitting data, as consistent annotations on out-of-sample data can not be reliably acquired. Similarly, recognizing tissue structures from unseen hematoxylin and eosin (H&E) stained pathology images is a widely-studied task often resolved in a supervised manner^33–36^. However, these tools tend to be more tissue type-specific and require substantial annotated training datasets.

To create a universal, robust, and generalizable tissue structure segmentation tool, we present a novel unsupervised partitioning method called Spatial Cellular Graph Partitioning (SCGP) in this work. SCGP is a fast and flexible method designed to identify the anatomical and functional units in human tissues. It can be effectively applied to both spatial proteomics and transcriptomics measurements. We further introduce a reference-query extension pipeline, SCGP-Extension, which enables the generalization usage of extending a set of reference tissue structures to previously unseen query samples. SCGP-Extension can address challenges ranging from experimental artifacts, batch effects, to disease condition differences and more, greatly enhancing SCGP’s robustness and versatility. To the best of our knowledge, SCGP is the first method validated on both spatial transcriptomics and proteomics data, as well as the first method purpose-built for generalization. We demonstrate its applications to five spatial proteomics and transcriptomics datasets collected on different tissue types comprising more than 1.6 million cells. The tissue structures identified by SCGP are evaluated against human annotations, benchmarked extensively against related software tools, and applied in downstream analysis that reveals disease-relevant biological insights.

## Results

### Unsupervised Partitioning of Spatial Cellular Graphs

To address the task of tissue structure identification, we developed a computational method SCGP that performs community detection on graph representations of tissue samples. Nodes in the graphs are small spatial units characterized by spatial coordinates and gene or protein expression at the location (**Methods**). In the representative case of multiplexed immunofluorescence (mIF) images^37^, nodes are defined on cells identified through the segmentation pipeline^20^(**Fig. 1A**). However, this concept of nodes can be further extended to accommodate broader spatial transcriptomics and proteomics data, such as spots in spatial transcriptomics sequencing measurements^7^ or small square patches in single-molecule fluorescence images^3^. In this study, we primarily present and discuss cell and spot based SCGP analysis. An alternative patch-based SCGP experiment can be found in **Supplementary Note 5** and **Supplementary Fig. 8**.

**Fig. 1.**
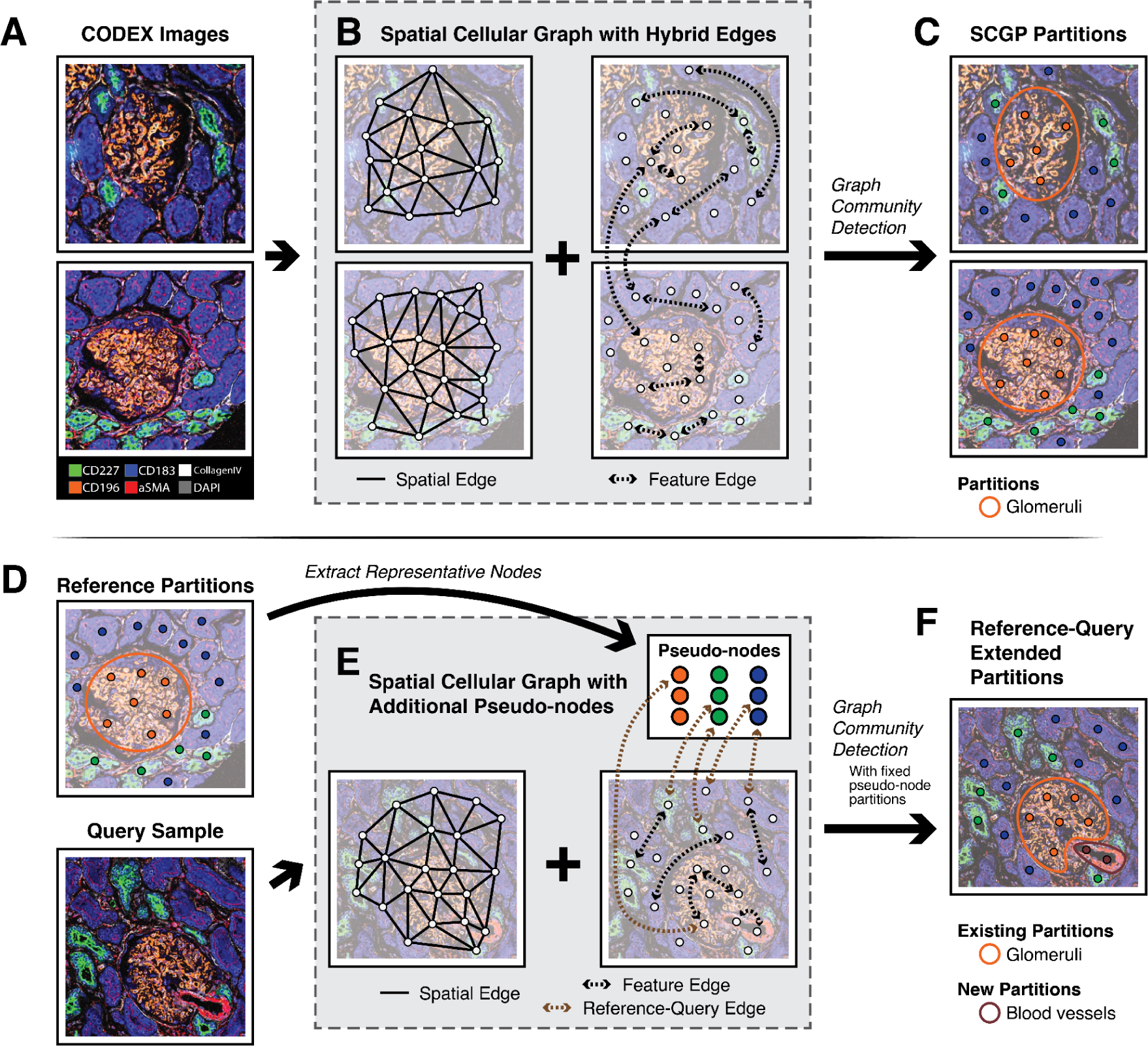
Workflow for SCGP and SCGP-Extension. **A.** Raw mIF images of example kidney samples show multiple tissue structures. **B.** Graph representations of mIF images are constructed on nodes (white circles) representing cells or other spatial units. Spatial edges (solid lines) and feature edges (dashed arrows) are constructed to reflect spatial closeness and feature similarity. **C.** Leiden graph community detection identifies partitions representing tissue structures. **D.** The query sample shares similar structures as the reference partitions. **E.** Graph representation of the query sample is constructed with additional pseudo-nodes (colored circles in the white box) extracted from the reference partitions. Reference-query edges (brown dashed arrows) are constructed between query nodes and pseudo-nodes. **F.** Leiden graph community detection yields both existing partitions that align with reference and new partitions that are previously unseen.

Two types of edges are constructed between nodes (**Fig. 1B** and **Methods**). Spatial edges are constructed between nodes based on Delaunay triangulation of node coordinates. These edges aim to capture the adjacency relationships between cells. Feature edges are constructed between nodes that share similar expression profiles.

Leiden graph community detection algorithm^38^ is then applied to the graphs, yielding partitions that represent the different tissue structures (**Fig. 1C**). The central aspect of this method is the joint contribution of two types of edges. Spatial edges guarantee the spatial continuity of the identified tissue structures, differentiating the method from cell type clustering in that multi-cell tissue structures will appear as cohesive entities. Feature edges interrelate tissue structures of the same type even if they are spatially separated (e.g., two glomeruli from different kidney samples), ensuring the consistency of tissue structure interpretation across samples. The relative quantity and contribution of each category of edge is a parameter that, while robust in our experiments, can be tuned. Throughout our experiments, we maintained a comparable amount of spatial and feature edges for community detection, see **Methods** for more details regarding hyperparameter settings of SCGP.

### SCGP Identifies Structures in Kidney Tissues

To examine the ability of SCGP to recognize known tissue structures, we assessed its performance on a cohort of 17 tissue sections from 12 individuals with diabetes and various stages of diabetic kidney disease (DKD)^39^. Tissue samples were imaged using the mIF platform CO-Detection by indexing (CODEX)^9^ and further annotated for four major kidney compartments: glomeruli, blood vessels, distal tubules, and proximal tubules. This cohort will be referred to as the DKD Kidney dataset (**Methods**) in the subsequent text.

Together with SCGP, we applied a diverse set of unsupervised annotation tools^13,15,24,27,31^ to the DKD Kidney dataset. All methods were applied to the combination of all 17 samples containing 137,654 cells, i.e., in a joint partitioning manner. Clustering/partitioning outputs on representative samples are visualized in **Fig. 2A** and **Supplementary Fig. 1A**, with the leftmost panels illustrating the raw mIF images with key biomarkers. Due to the unsupervised nature of the output, we reordered the output clusters of each method in accordance with manually annotated compartments. In **Fig. 2A**, the top panel for each column shows the output, and the bottom panel highlights the mismatches. Across samples, partitions annotated by SCGP frequently demonstrated the highest fidelity to manual annotations.

**Figure 2.**
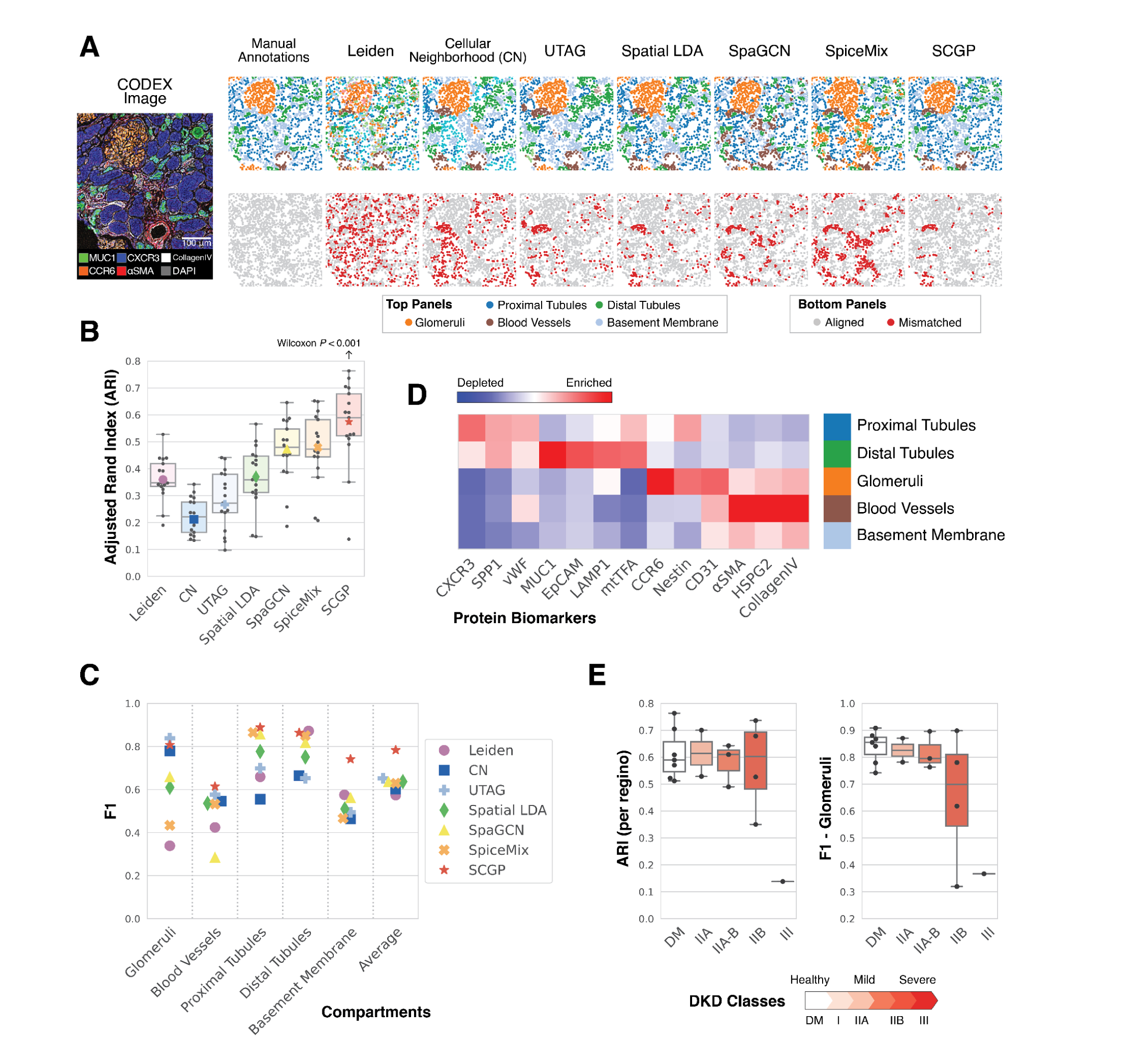
Unsupervised annotations of DKD Kidney samples. **A.** Annotations from SCGP and other unsupervised annotation methods recognize tissue structures aligned with manually annotated compartments. Nodes representing cells are colored according to the assigned clusters/partitions in the top panels, colors not listed in the legend (e.g., cyan) refer to clusters/partitions that cannot be matched to any compartment. Mismatched nodes are highlighted in red in the bottom panels. **B.** ARIs are calculated between unsupervised annotations and manual annotations. SCGP performed significantly better than all other methods (*P* < .001, Wilcoxon signed-rank test). **C.** For each manually annotated compartment, F1s are calculated between manual annotations and the most overlapped cluster/partition. **D.** Signature protein biomarkers for SCGP-identified partitions match expectations of kidney tissue structures. **E.** SCGP annotations on samples with different classes of DKD show varying levels of alignment accuracy. DM (i.e., healthy kidney) and DKD classes represent the different progression stages, assigned by following the Tervaert classification^40^.

We calculated Adjusted Rand Index (ARI) between manually annotated compartments and the unsupervised partitions to evaluate their alignments (Fig. 2B). SCGP achieved the top performance with a mean ARI of 0.57 (SD = 0.15), significantly outperforming all other methods (*P* < .001, Wilcoxon signed-rank test). Notably, ARI downweighs compartments that are smaller in size, while glomeruli, though having less coverage, are the major functional units in kidneys and carry much greater importance in downstream analysis. To break down the performance metrics, we calculated F1 scores for individual compartments between manual annotations and the most overlapped cluster/partition (**Methods** and Fig. 2C). We found that UTAG^15^, Cellular Neighborhood (CN)^13,26^, and SCGP performed the best at recognizing glomeruli (F1 ∼ 0.8), while SpiceMix^24^, SpaGCN^31^, and Spatial LDA^27^ excelled at recognizing tubule structures. Overall, SCGP achieved best average accuracy in discerning all the manually annotated compartments.

We further looked into the details of the SCGP-identified partitions by highlighting signature protein biomarkers that are enriched in each partition (Fig. 2D and **Supplementary Fig. 1B**). The heatmap corresponds well to our expectation, with CCR6^41^ and Nestin^42^ among the top biomarkers for glomeruli, CXCR3^43^ and MUC1^44^ for proximal and distal tubules. Interestingly, by grouping performances based on the disease progression^40^, we found that unsupervised annotations aligned better with manually annotated compartments in healthy (DM) or mild DKD samples (DKD Class I, IIA). The substantial decrease in ARI and F1 for severe DKD samples (DKD class IIB, III) indicates how normal tissue structures and functions are dysregulated in DKD (Fig. 2E).

### SCGP Identifies Layers in Human Brain Data

Next, we assessed SCGP’s performance on a spatial transcriptomics dataset from human dorsolateral prefrontal cortex (DLPFC)^45^ that was acquired using the Visium platform^7^. In contrast to the mIF approach, the Visium platform features a grid of spatially barcoded oligonucleotide arrays that can be used for mRNA capture and library preparation. Each array (i.e., spot) consists of multiple cells. We adapted our method by treating each spot as a node and defining edges based on the grid and gene count information (**Methods**).

DLPFC contains 12 individual samples annotated with 7 compartments: 6 cortical layers (L1-6) and white matter (WM). We directly compared our method against existing tools developed for spatial transcriptomics data, including BayesSpace^28^, SpaGCN, and SpiceMix, in recognizing manually annotated layers. In the first experiment, we conducted clustering/partitioning on each sample independently. Results on a representative sample are demonstrated in Fig. 3A, revealing a clear layer-wise pattern in the tissue structures. Among the benchmarked methods, SCGP and SpiceMix achieved top ARI scores, exhibiting superior alignment with the ground truth compartments marked by dashed lines. We calculated quantitative metrics for all 12 samples (**Fig. 3B-C**) and demonstrated that SCGP achieves comparable, if not superior, performance (median ARI = 0.51, median F1 = 0.65) as the unsupervised annotation tools designed for spatial transcriptomics.

**Figure 3.**
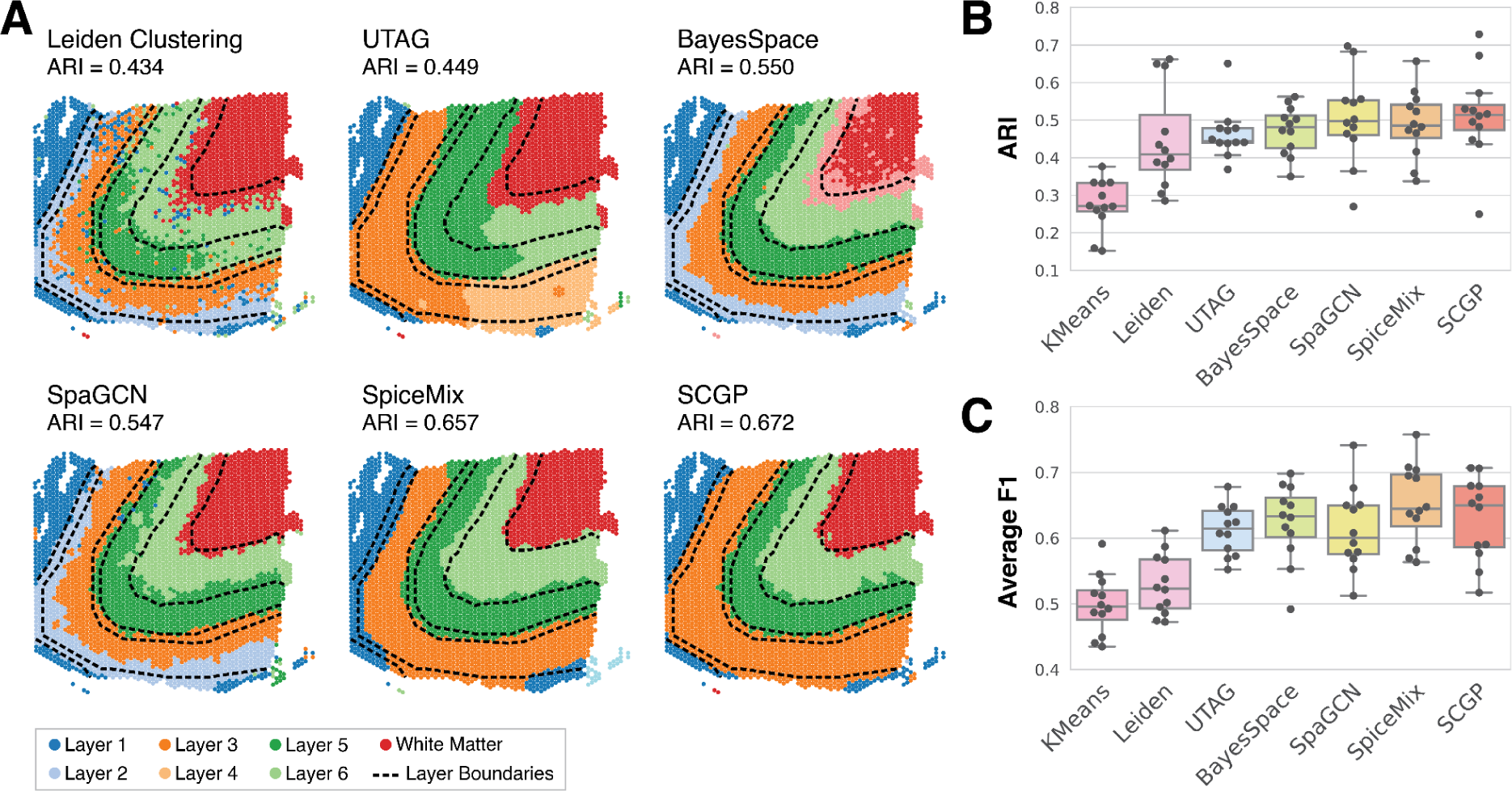
Unsupervised annotations of human DLPFC samples. **A.** Annotations from SCGP and other benchmarked methods on a representative sample (151673). Note that Layer 2 and Layer 4 are not fully recognized. Boundaries between ground truth layers are annotated as dashed lines. **B.** ARIs are calculated between unsupervised annotations and ground truth layers on 12 samples, with each sample annotated independently. **C.** F1s are calculated on each layer between unsupervised annotations and ground truth. Average F1s for all 12 samples are illustrated in the box plot.

We then evaluated if joint partitioning the combination of multiple samples can help improve performances. Following the experiment design by Chidester et al.^24^, we applied unsupervised annotation tools to the combination of four samples collected from the same donor (**Supplementary Fig. 2A, C, D**). Methods including BayesSpace, SpaGCN, and SpiceMix performed better than independent clustering results, while UTAG and SCGP yielded worse results. To address the performance drop, we adopted the solution described in the next section: extending the superior partitions from the selected reference sample to the remaining samples.

### SCGP Extends Existing Tissue Structures to Unseen Samples

In practical applications, it is often advantageous to conduct the initial clustering/partitioning on a selected subset of high-quality samples, inspect them for validity, then extend the resulting annotations to a wider range of data. This generalization approach can be useful for performing inference on prospective data, suppressing unwanted noise or batch effects, comparing samples across disease conditions, and detecting unseen disease states. However, existing unsupervised annotation tools have very limited support for this functionality. Most methods (e.g., UTAG, SpaGCN) require either retraining or re-fitting the clustering model or the addition of separate prediction models to extend existing partitions. To address this need, here we present a specialized reference-query extension pipeline SCGP-Extension to effectively handle the generalization usage.

SCGP-Extension begins with partitioning a group of high-quality reference samples. Resulting partitions are assumed to represent the ground truth structures and are referred to as reference partitions (Fig. 1D). Through random sampling of nodes from the reference data, we construct representative nodes for each reference partition, named “pseudo-nodes” (**Methods**). Next, given unseen query samples, their graphs are constructed with explicit addition of the pseudo-nodes (Fig. 1E), which connect to query nodes based on feature similarity. The following graph community detection step is conducted with partitions of the pseudo-nodes pre-assigned and fixed. Consequently, query nodes resembling pseudo-nodes are assigned the corresponding partition. Query nodes that do not resemble any reference partitions will form their own groups and be assigned as newly discovered partitions (Fig. 1F).

To demonstrate how SCGP-Extension improves partitioning performance, we revisited the joint partitioning experiment of DLPFC samples. Provided that independent partitioning on one representative sample yielded successful results (Fig. 3A), we adapted this sample as a reference and extended its partitions to the rest. SCGP-Extension outputs considerably outperformed SCGP (**Supplementary Fig. 2A**), achieving higher alignment (ARI) and accuracy (F1) scores (**Supplementary Fig. 2C-D**) comparable with other spatial transcriptomics annotation methods. Additionally, we tested methods that utilize partial ground truth, in which the extension of labels outperformed predictive modeling (**Supplementary Fig. 2B** and **Supplementary Note 1**).

Moreover, SCGP-Extension can generalize across disease conditions and help identify unseen disease states. Without the extension pipeline, we observed suboptimal SCGP partitions in severe DKD samples from the DKD Kidney dataset (**Fig. 4A, Fig. 2E** and **Supplementary Fig. 1A**), where glomeruli exhibited elevated levels of fibrosis (e.g., Collagen IV) and decreased expression of native biomarkers (e.g., CCR6). When jointly partitioning these samples with healthy and mild DKD samples, fibrotic glomeruli were often misrecognized as blood vessels (black arrows in **Fig. 4A**). Similarly, predictive models trained with manual annotations showed limited success in identifying fibrotic glomeruli (**Fig. 4A**, XGB Prediction). To overcome these challenges, we applied SCGP-Extension to generalize primary partitions from healthy and mild DKD samples to severe DKD samples. SCGP-extension yielded superior results (**Fig. 4A**, SCGP-Extension), preserving most of the original partitions while uncovering two new structures: the purple partition outlined fibrotic glomeruli, characterized by the depleted native biomarkers and enriched Collagen expression (**Fig. 4B**); and the red partition, characterized by elevated CD45 and CD68 expression (**Fig. 4B**), suggesting the infiltration of immune cells (e.g., macrophages). Compared to the joint partitioning and predictive model outcomes, SCGP-extension delivered more accurate results both visually and quantitatively (**Fig. 4C**). Furthermore, it highlighted partitions of unseen disease states, offering valuable insights into the process of disease progression.

**Figure 4.**
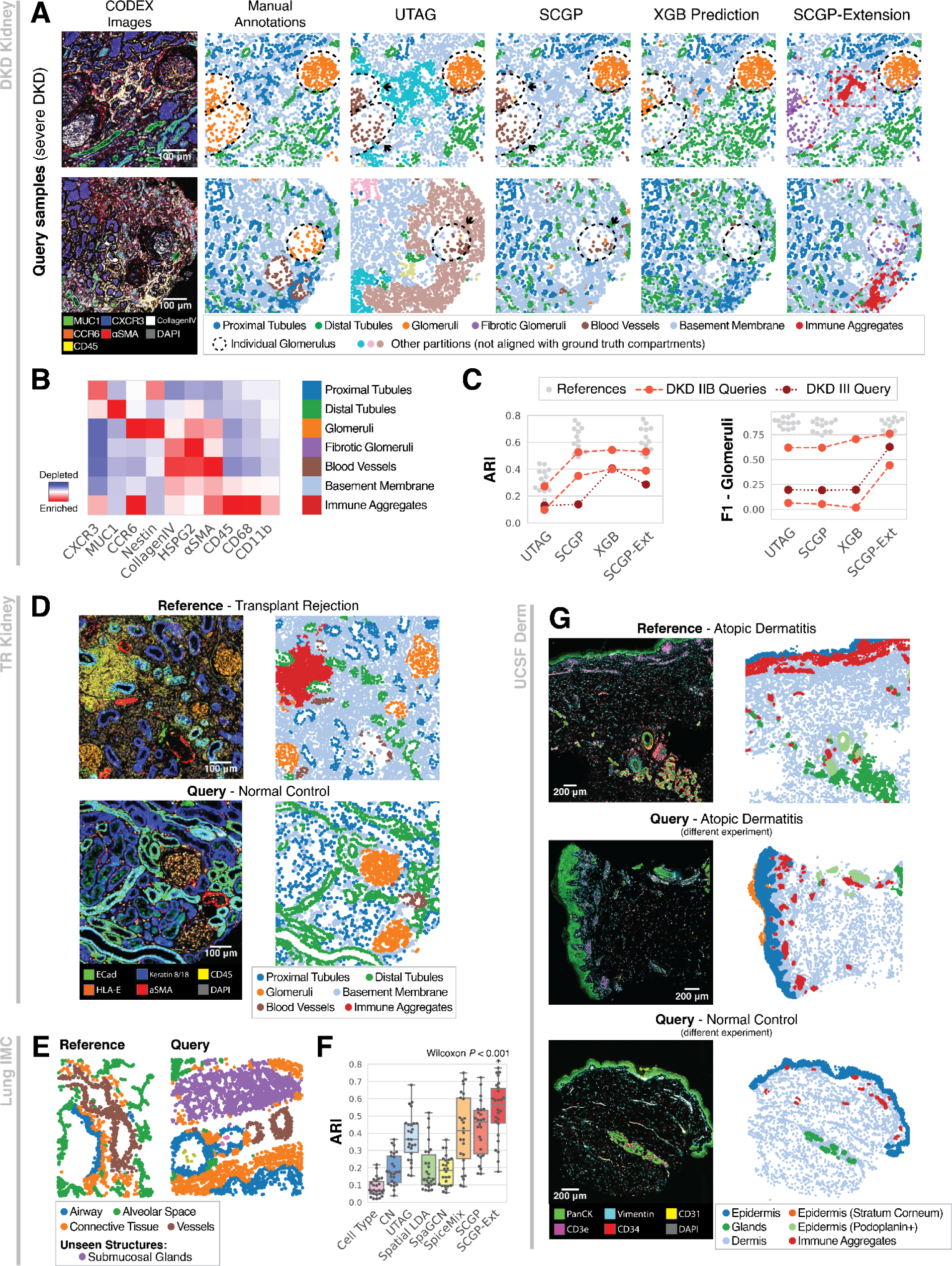
Versatile applications of SCGP-Extension. **A.** Compared to joint partitioning or predictive models, SCGP-Extension better recognized the fibrotic glomeruli (purple dashed circles) in severe DKD samples, and identified a previously unseen immune aggregates partition (red boxes). **B.** Extension to severe DKD samples improved partition alignment and accuracy. **C.** Heatmap shows signature protein biomarkers for extended partitions in the severe DKD samples, note the additional fibrotic glomeruli and immune aggregates partitions. **D.** In the TR Kidney dataset, SCGP partitioned a cohort of kidney samples with heavy inflammation (top row). SCGP-Extension further extended the partitions to control samples with minimal immune responses (bottom row). **E.** In the Lung IMC dataset, SCGP-Extension recognized a new tissue structure submucosal glands (purple) in the query samples that is unseen in references. **F.** SCGP and SCGP-Extension aligned better with manual annotations of lung samples. SCGP-Extension achieved significantly better ARIs than all other methods (*P* < .001, Wilcoxon signed-rank test). **G.** In the UCSF Derm dataset, SCGP-Extension consistently partitioned samples from multiple experiments with different skin conditions.

We further assessed the performance of SCGP on a separate dataset comprising kidney tissue samples from patients who experienced transplant rejection (TR Kidney dataset, **Methods**). The analysis derived the same set of tissue structures from samples regardless of differences in disease conditions and biomarkers (**Fig. 4D**). Furthermore, to compare samples exhibiting various degrees of immune responses, we extended partitions from reference samples with heavy inflammation and immune signatures to query samples with minimal inflammation. SCGP-Extension consistently annotated structures of the same type (**Fig. 4D** and **Supplementary Fig. 3D**), highlighting the different distributions of tissue structures in these two conditions (**Supplementary Note 2**). Notably, SCGP-Extension effectively dealt with a variable background artifact present in one sample of the TR Kidney dataset (**Supplementary Fig. 6**).

SCGP-Extension can identify unseen anatomical structures in query samples. To assess the broader applicability of SCGP and SCGP-Extension, we evaluated our methods on an Imaging Mass Cytometry (IMC) dataset collected on healthy lung specimens (Lung IMC dataset, **Methods**), which also contains manual annotations for anatomical structures. SCGP achieved remarkable alignment with manual annotations (median ARI=0.464), but tissue structures across samples were assigned to different partitions (**Supplementary Fig. 4A**). To address this issue, we conducted primary SCGP on a well-integrated subset of samples and extended partitions to the remaining samples (**Supplementary Note 3**). Notably, SCGP successfully identified two anatomical structures (submucosal glands and cartilage) that were absent in the reference, assigning them as unseen tissue structures while preserving all the known structures (Fig. 4E). SCGP-Extension also achieved significantly better alignment scores (median ARI=0.564, Fig. 4F and **Supplementary Fig. 4B-C**) than all other benchmarked methods (*P* < .001, Wilcoxon signed-rank test).

SCGP-Extension can also help mitigate batch effects between experiments. On a cohort of skin samples collected from four separate experiments (UCSF Derm dataset, **Methods**), we employed different unsupervised annotation methods to define tissue structures. The UCSF Derm dataset comprised samples of different skin conditions, sharing similar anatomical structures (**Supplementary Table 1**). Both UTAG and SCGP failed to link tissue structures from different samples (**Supplementary Note 4** and **Supplementary Fig. 7A-B**). We then attempted SCGP-Extension by defining reference partitions on samples from one experiment (**Supplementary Fig. 5A**) and extending them to the rest (**Supplementary Fig. 5B-D**). Regardless of batch effects between experiments and differences in disease conditions, SCGP-Extension successfully recognized consistent partitions reflecting anatomical structures across samples (Fig. 4G, **Supplementary Fig. 7D**).

### SCGP Partitions Assist Downstream Analysis of Disease States

Partitions acquired by unsupervised annotation using SCGP reflect the anatomical and functional structures of the subject tissues. In this section we use two datasets to demonstrate how these tissue structures can facilitate analysis of disease states and enable the discovery of biological insights regarding disease-relevant partitions.

In the DKD Kidney dataset, samples were collected from individuals with different DKD classes. Based on the partitioning of these samples, we were interested in exploring correlations between tissue structures and disease progression. Fig. 5A illustrates three representative samples of different DKD classes along with their tissue structures annotated by SCGP and SCGP-Extension. Clear visual differences between samples can be observed: Tubules and glomeruli were much denser in the healthy sample, while these structures gradually deformed over the course of DKD, accompanied by fibrosis and infiltration of immune cells. These changes were also reflected in the tissue structures: A significant increase (*P* < 0.001, Jonckheere-Terpstra test^46^) in area proportion of the basement membrane partition was observed across samples (Fig. 5B), reflecting the degradation of normal kidney structures. An immune aggregate partition (red) and a fibrotic glomerular partition (purple) were identified in the severe DKD sample with SCGP-Extension, which were not present in healthy and mild DKD samples.

**Figure 5.**
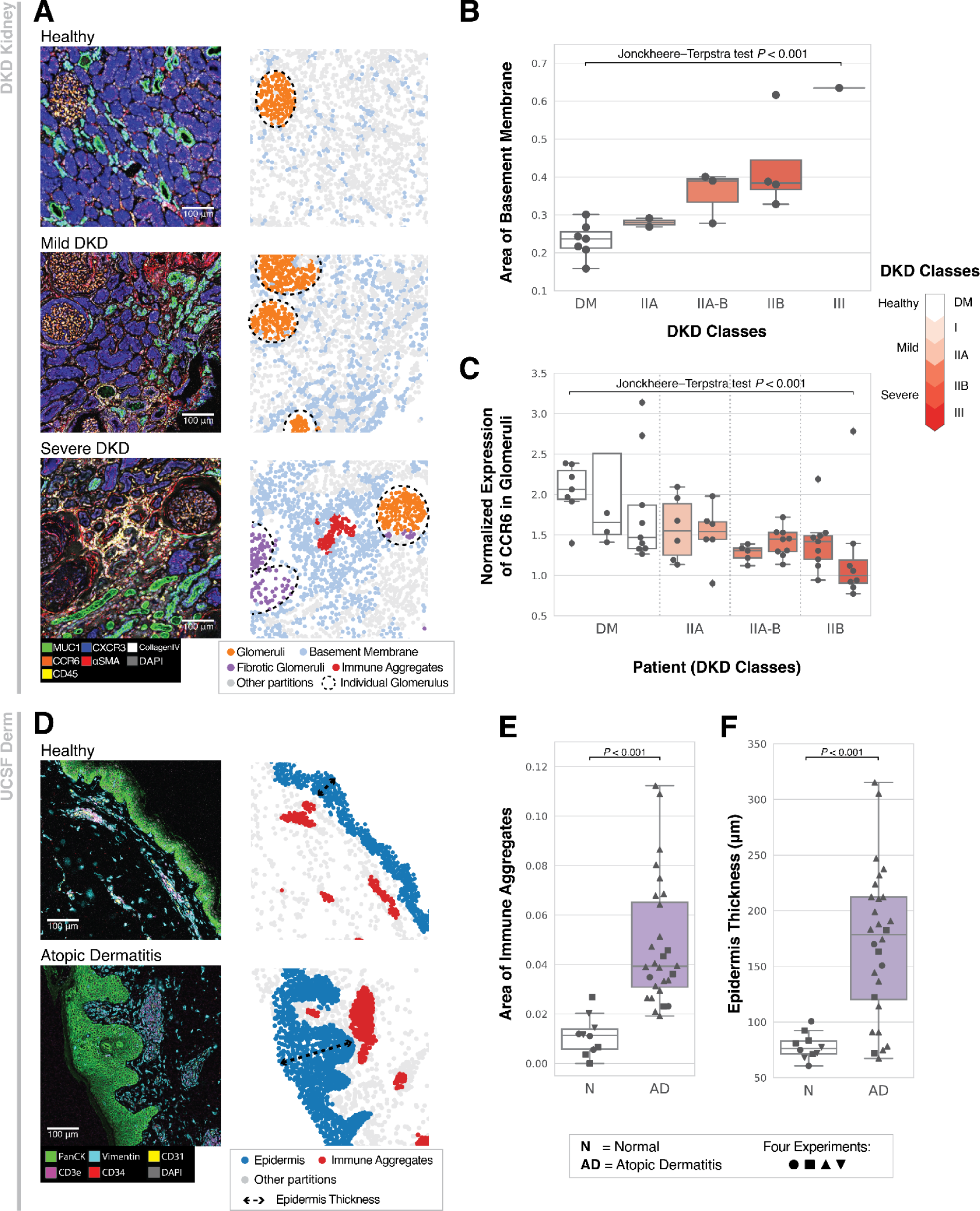
Downstream analysis of disease states with SCGP-identified partitions. **A.** Three representative samples of different DKD classes are illustrated, note the fibrosis of glomeruli and increase in the area of the basement membrane partition (light blue-colored nodes). Glomeruli are segmented by dashed circles. **B.** Box plot shows that the area proportion of the basement membrane partition significantly increases (*P* < 0.001, Jonckheere-Terpstra test) in DKD samples. Each dot represents a tissue sample. **C.** Expression of native proteins (CCR6) significantly decreases (*P* < 0.001, Jonckheere-Terpstra test) in glomeruli of DKD patients. Each dot represents an individual glomerulus, and each box summarizes glomeruli from one patient. Note the heterogeneity observed within a single tissue sample (**A.**, bottom row) and between patients with the same DKD class. **D.** Two samples from the UCSF Derm dataset with different skin conditions are illustrated. **E.** Area proportion of the immune aggregates partition significantly increases in atopic dermatitis samples (*P* < 0.001, two-sided two-sample t-test). **F.** Epidermal layers show significant thickening in atopic dermatitis samples (*P* < 0.001, two-sided two-sample t-test).

Furthermore, to characterize how DKD affects glomerular functions, we assessed the protein biomarker expression of individual glomerulus (dashed circles in Fig. 5A), annotated by deriving connected components of the SCGP glomeruli partitions. Results suggested that glomeruli undergo significant loss of native proteins (CCR6) throughout the disease progression (*P* < 0.001, Jonckheere-Terpstra test, Fig. 5C), with high intra-sample and inter-sample heterogeneity.

We next applied the partition-based analysis to the UCSF Derm dataset comprising normal samples and atopic dermatitis samples. Representative samples visualized in Fig. 5D shows notable differences in terms of epidermal thickness and immune cell densities. We verified these visual signatures using SCGP partition information: Area proportion of the immune aggregates partition was calculated and demonstrated significant increases in atopic dermatitis samples (*P* < 0.001, two-sided two-sample t-test, Fig. 5E). Thickness of the epidermal layer was characterized using the contour of the epidermis partition, which showed significant increases as well (*P* < 0.001, two-sided two-sample t-test, Fig. 5F). Patients with atopic dermatitis also exhibit a much more heterogeneous distribution of epidermal thickness^47^.

## Discussion

In this work, we present SCGP, an unsupervised annotation tool for spatial transcriptomics and proteomics measurements. SCGP embeds spatial information and biomarker expression of subject tissues into graph representations, and performs Leiden graph community detection to identify partitions corresponding to anatomical and functional structures. The reference-query extension pipeline, SCGP-Extension, further boosts the potential of the method by enabling generalization of existing tissue structures to unseen samples. Our experiments demonstrate the power of SCGP and SCGP-Extension in identifying tissue structures in various data cohorts and show how tissue structures assist downstream biomedical research and discoveries.

We compared SCGP against a list of unsupervised annotation tools that similarly utilize both spatial information and molecular profiling output. Representative methods such as CN, UTAG, and Spatial LDA define different concepts of neighborhoods, usually based on distance thresholding, and annotate them through unsupervised clustering. These methods tend to work well only for structures of specific spatial scales. Another class of methods use graph representations of tissue samples, where the spatial organizations of cells are modeled in the graph structures. Computational tools including latent variable modeling and graph neural networks are applied to annotate nodes (cells) according to their graph context, but a large drawback is the separation between spatially disconnected samples. SCGP follows the graph representation approach and enhances it with nearest neighbor feature edges, thereby weaving all samples into one cohesive graph.

Moreover, SCGP-Extension is the first method that addresses the long-standing need of generalizing structures to previously unseen samples. SCGP-Extension resembles supervised learning tools in that models apply knowledge learned from the reference samples (i.e., training data) to unseen query samples (i.e., test data). Simultaneously, SCGP-Extension can isolate novel tissue structures during inference. In practice,we have demonstrated that SCGP-Extension can help overcome common challenges including experimental artifacts, batch effects, and different disease conditions. Additionally, it can help uncover unseen disease states and anatomical structures.

SCGP and SCGP-Extension offer outstanding running time performance compared to many unsupervised annotation tools, some of which require extensive parameter estimations or optimizations (**Supplementary Note 6** and **Supplementary Table 2**). In practice, this advantage allows for applications to much larger datasets and facilitates model parameter tuning to obtain optimal partitions.

SCGP and SCGP-Extension are validated to manage various spatial proteomics and transcriptomics measurements effectively, requiring only minimal parameter adjustments (**Methods**). The two major parameters involved are the density of feature edges and the granularity of partitioning. In our experiments, both were maintained in similar settings, with a comparable number of feature edges and spatial edges, and resolution parameters that yield five to six partitions. Specifically for SCGP-Extension, an additional parameter, the extent of extension, controls the balance between generalizing existing partitions and exploring new partitions. In examples involving different disease conditions, certain tissue structures may experience changes in their expression profiles (e.g., fibrosis, immune infiltration), and the decision of whether to integrate or separate these structures will depend on the specific downstream applications.

Nevertheless, we acknowledge the presence of certain caveats in SCGP and SCGP-Extension, which hinder their applications in specific scenarios. One limitation is that SCGP can only reliably detect coarse structures (i.e., four-seven structures), especially during joint partitioning of multiple samples. Increasing granularity will cause structures of the same type from different samples being assigned to separate partitions, complicating interpretations. The other disadvantage is that our methods appear to be less suitable for identifying thin-layer structures. Due to the design of the hybrid graph, spatial edges are isotropic for the purpose of community detection. Thin-layer structures exhibiting much denser spatial connections in their normal directions than tangential directions are harder to detect. In the DLPFC study, alternative methods (e.g., BayesSpace, SpiceMix) can identify thinner structures under more granular settings, while SCGP tends to bisect existing layers in the orthogonal direction. Refining the spatial edges to reflect the anisotropy of tissues would be a direction to improve the performances of SCGP.

In addition to addressing the aforementioned drawbacks, several avenues for improving SCGP can be pursued in future research. One direction is the integration of multi-modal data, such as morphology embeddings from H&E staining, which would greatly enrich the feature space and allow SCGP to account for cellular and tissue-level morphological differences. Secondly, rather than performing fully unsupervised partitioning from scratch, incorporating prior knowledge of the expected tissue structures to SCGP, similar to how reference partitions contribute to SCGP-Extension, might yield better-aligned results. Looking ahead, SCGP opens new opportunities for analyzing and understanding spatially-resolved molecular profiling data. By inserting a middle layer between cell-level annotations and sample/patient-level characteristics, it facilitates research and discoveries by enabling better dissection of samples and encouraging analysis tailored to specific structures-of-interest.

## Methods

### Datasets

We used five spatial proteomics and transcriptomics datasets collected from diverse tissue types in this work. An overview of the statistics and primary phenotypes can be found in **Supplementary Table 1**. This subsection will provide additional information on each dataset. Please refer to the source publications for full details.

### DKD Kidney Dataset^39^

Kidney samples were obtained from patients with diabetes and healthy kidneys (DM, five individuals), DKD classes IIA, and IIB (two individuals per class), IIA-B intermediate (two individuals), and III (one individual). Twenty-three cores, each 0.5 mm in diameter, were sampled from twelve tissue blocks and assembled into a tissue microarray (TMA). The TMA block was further sectioned into 5 µm slices.

A TMA section was imaged and characterized using the CO-Detection by indexing (CODEX) platform. After excluding medulla samples and quality control, a total of 17 cortical section samples across various DKD classes were acquired. Each sample was imaged for 21 protein biomarkers (see columns in **Supplementary Fig. 1B**).

### DLPFC Dataset^45^

Spatial gene expression in human postmortem DLPFC tissue sections was profiled using two pairs of “spatial replicates” from three independent neurotypical adult donors on the Visium platform, each pair comprising two directly adjacent, 10-µm serial tissue sections, with the second pair located 300 µm posterior to the first. In total 12 samples are collected and examined. We downloaded the filtered count matrices for all 12 samples from the spatialLIBD project^48^.

In the independent partitioning experiment (Fig. 3), we filtered the count matrices to exclude spike-in genes, mitochondrial genes, and genes that have nonzero expression in fewer than three spots. Expression matrices were normalized to have the same total counts per spot (median of all pre-normalize spots), log-transformed and reduced to the top 50 principal components.

In the joint partitioning experiment (**Supplementary Fig. 2**), we followed the preprocessing steps outlined in SpiceMix^24^: Genes having nonzero expression in less than 10% of spots were removed. Expression matrices were normalized to have total counts of 10,000 per spot and log-transformed. We further reduced the expression to the top 40 principal components.

### TR Kidney Dataset

Kidney samples were obtained from patients who underwent allograft nephrectomy, as previously described^49^. Briefly, a TMA was constructed using 2 mm cores of cortical tissue. The TMA comprised 7 samples of normal, peritumoral renal cortex (from patients undergoing native nephrectomy for tumor removal), and 43 samples of cortex from patients undergoing allograft nephrectomies. After sectioning of the tissue block, only 41 cores from allograft nephrectomies remained.

An FFPE embedded tissue microarray core of human kidney samples were stained and acquired on the PhenoCycler Fusion using a 51-plex biomarker panel (**Supplementary Fig. 3C**) by the Enable Lab. 5 normal and 17 transplant rejection kidney samples were used in this study.

### Lung IMC Dataset^15,50^

Lung samples were acquired from three healthy donor lung specimens. In total 26 samples were imaged using IMC with 28 biomarkers (see columns in **Supplementary Fig. 4D**). Tissue samples were collected with a particular focus on airways extending from proximal bronchi and succeeding divisions to terminal and respiratory bronchioles.

Each image was manually annotated with organ-specific microanatomical domains: airways, connective tissue, submucosal glands, vessels, cartilages, and alveolar space. These manually annotated domains were used as labels for unsupervised annotations. Additionally, cells in these samples were phenotyped into seven broad clusters of cell identity: CD8 T cells, macrophages, mast cells, smooth muscle cells, endothelial cells, epithelial cells, and connective tissue cells. Cell type information was used in CN and Spatial LDA, other methods only used biomarker expression data.

We downloaded preprocessed biomarker expression matrices and domain/cell type annotations from the source publication^51^.

### UCSF Derm Dataset

Tissue samples were acquired as 2-4 mm punch or shave skin biopsy specimens from healthy control and atopic dermatitis patients. In total, 44 skin samples were obtained and stained in 4 experiments with varying biomarker panels by the Enable Medicine Lab. See **Supplementary Table 1** for a detailed breakdown of experiments and patient skin disorders. 35 shared protein biomarkers were used in the unsupervised annotation analysis with UTAG and SCGP.

### Preprocessing

The Visium DLPFC dataset was downloaded and normalized as specified above, with no additional processing or batch correction executed. The Lung IMC dataset was downloaded as matrices of preprocessed and normalized biomarker expression, no additional processing was executed.

For the three CODEX datasets, we followed the preprocessing pipeline established in the prior work by Wu et al.^16^ Briefly, a neural network-based cell segmentation tool DeepCell^20^ was applied to DAPI images to identify nuclei, which were further dilated to obtain whole-cell segmentation.

Next, the biomarker expression for biomarker *j* in cell *i* was computed following the strategy below^52^:

- For channel *j*, mean pixel intensity within the cell segmentation mask of cell *i* was calculated and denoted as 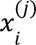. The set of expression values for all cells in the same sample {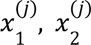, …} was denoted as *X*^(*j*)^.
- Normalized expression value for channel *j* was calculated using quantile normalization and arcsinh transformation:

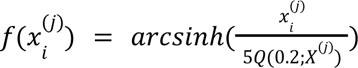 in which *Q*(0. 2; *X*^(*j*)^) represents the 20-th quantile of *X*^(*j*)^ and *arcsinh* is the inverse hyperbolic sine function. The set of all normalized expression values {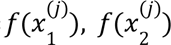, …} was denoted as *f*(*X*^(*j*)^.
- z-score of normalized expression value was calculated:

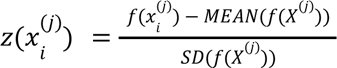 It should be noted that SCGP does not require any cell clustering or classification inputs, it infers partitions using biomarker expression values of cells.

### Spatial Cellular Graph Partitioning (SCGP)

SCGP is an unsupervised annotation tool that recognizes tissue structures by partitioning graphs constructed based on the spatial organization of cells (or other units) in the subject tissue sample(s). The SCGP pipeline comprises the following steps.

### Construction of Nodes

Nodes represent small spatial regions in the tissue, and are indivisible units throughout the partitioning process. In this study, we employed three different strategies for defining nodes:

- Cells: Nodes are defined based on individual cells, which are identified via the cell segmentation preprocessing step specified above. Biomarker expression values are calculated and normalized accordingly, and these values are set as the node features.
- (Visium) Spots: In the DLPFC dataset, nodes are defined based on the barcoded spots used in the Visium platform, each measuring gene expression in a circular area 55 µm in diameter. The normalized gene expression values or top principal components are set as node features.
- Patches: In the patch-based SCGP experiment (**Supplementary Note 4**), nodes are defined based on small square patches with 12 µm side lengths on the mIF images, sampled using a sliding window mechanism (stride equals patch side length). Only patches within the spatial range of tissue are retained (>85% overlap), and the average fluorescence intensities for each biomarker channel are used as node features.

### Construction of Spatial Edges

Spatial edges are constructed between spatially adjacent nodes to embed the spatial closeness relationship into the graph.

- Cells: A Delaunay triangulation is conducted on centroid coordinates of all cells from the same sample. Node pairs that share edges in the triangulation output are connected, excluding any edge exceeding 35 µm in length.
- (Visium) Spots: Nodes (i.e., spots) are spatially arranged in a close-packing manner. Each node is connected to its six closest neighboring nodes.
- Patches: Nodes (i.e., patches) are spatially arranged in a regular 2D grid. Each node is connected to its four immediately adjacent nodes.

### Construction of Feature Edges

Feature edges are constructed between nodes with similar biomarker expression profiles. For node *i*, its *k* nearest neighboring nodes in the expression space are identified based on Euclidean distances between node features (e.g., z-scored protein expression values, principal components of the gene expression). We used the nearest neighbor descent^53^ approximate queries implemented in `PyNNDescent` for better computational efficiency.

In practice, *k* is a hyperparameter that controls the balance between spatial coherence and expression consistency within partitions. We typically set it to an integer that delivers a similar amount of feature edges as spatial edges, usually between 3 and 6. Minor changes in *k* do not change partition outcomes in our experiments, but a larger *k* (*k* > 8) might result in spatially fragmented partitions.

### Graph Community Detection

Nodes, spatial edges, and feature edges define the spatial cellular graph input for SCGP. We used the Leiden algorithm^38^ to detect graph communities.

We adapted the python implementation in `leidenalg`, and we used the Constant Potts Model^54^ as the quality function for community detection (`leidenalg.CPMVertexPartition`). Additional arguments include:

- Edge weights: For each edge, its weight is defined as the inverse of Euclidean distance between the node features of the two nodes it connects. We further normalized all edge weights by their median value.
- Resolution parameter (γ): γ controls the density of the output communities.

Note that the resolution parameter γ is the second major hyperparameter of SCGP, regulating the granularity of the output partitions. We empirically tuned γ to generate 4-7 partitions based on our understanding of the corresponding tissues. Higher γ will sometimes result in the same tissue structures being assigned to different partitions in different samples. This is likely due to the fact that inter-sample feature edges are sparser (and hold less weights) than intra-sample edges.

### Post-processing

Upon acquiring the initial partition outcomes, optional post-processing steps can be executed to refine the results:

- Size Filtering: Partitions accounting for less than 0.2% of all nodes are discarded.
- Spatial smoothing: For any node who holds a different partition assignment from all its spatial neighbors, we reassigned it to the partition held by the majority of its spatial neighbors (>50%) if applicable. This step will remove the rare cases where a lone node appears in a spatially-coherent structure.

### SCGP-Extension (Reference-Query Extension)

SCGP-Extension extends a given set of reference partitions to unseen query samples. Query samples are processed in the same manner as specified in the SCGP pipeline for graph construction. Reference partitions are defined on the reference nodes, which should be in the same format as nodes in the query samples. These partitions are usually generated through a primary SCGP run on the reference samples. It is worth noting that technically any form of discrete labels can be employed as reference partitions. **Supplementary Note 1** demonstrates an example experiment that extended ground truth labels to query samples.

### Construction of Pseudo-nodes

The major difference between SCGP-Extension and SCGP is the introduction of pseudo-nodes that serve as guidance for query sample partitioning. Based on the reference nodes and their reference partitions, pseudo-nodes can be created for each partition via two strategies:

- Selection: A median node feature vector is calculated using all reference nodes affiliated with the partition. The Euclidean distance between each node’s feature vector and the median vector is calculated. Representative nodes that have closest distances to the median vector are selected as pseudo-nodes.
- Random sampling: The mean and covariance of the node feature vector are calculated using all reference nodes affiliated with the partition. Pseudo-nodes are generated by sampling multivariate normal random variables based on the mean and covariance.

In practice, we typically generate 100 pseudo-nodes for each partition, but the size can be adjusted based on the number of nodes in the reference and query samples. The two strategies tend to yield comparable results, we empirically prefer the selection strategy.

Note that prior to their integration into the query graph, dense feature edges are added to the pseudo-nodes – 20 nearest neighbors for each pseudo-node if in total 100 nodes per partition are used. This is to guarantee well-structured communities within the pseudo-nodes.

### Construction of Reference-Query Edges

Pseudo-nodes are then integrated into the query graph via additional reference-query edges. For each node in the query graph, its *k*’ nearest neighbors in the pseudo-nodes based on Euclidean distances between node features are identified. An additional ratio parameter *r* (0 < *r* ≤ 1) is included to regulate the strengths of reference-query edges. Given a total of *N* nodes in the query graph, the initial nearest neighbor search will yield *Nk*’ reference-query edges. These edges are sorted by edge weights (i.e., Euclidean distances), with only the top *rNk*’ edges with the highest weights or closest distances retained.

The completed query graph comprises spatial edges, intra-sample feature edges (*k*-nearest neighbors within the query nodes), and reference-query feature edges (*r*-downsampled *k*’-nearest neighbors between query nodes and pseudo-nodes). *r* and *k*’ are two additional hyperparameters in SCGP-Extension, which control the degree of matching between query nodes and reference partitions. In practice we adjusted *k*, *k*’ and *r* so that the total number of feature edges slightly surpasses the number of spatial edges (∼1.2x), with an equal number of intra-sample feature edges and reference-query feature edges. *r* is typically set to 0.5, which ensures that nodes from unseen tissue structures are not excessively connected to the pseudo-nodes, thereby avoiding forced alignment with existing reference partitions. However, based on the task and understanding of subject tissues, *r* can be set to a higher value for enhanced alignment.

### Graph Community Detection with Fixed Membership Assignment

The same community detection strategy is used to identify partitions in the query sample: Leiden algorithm with the Constant Potts Model as quality function. Additional arguments including edge weights and resolution parameter γ are specified in the same manner as specified in the SCGP pipeline.

Notably, as pseudo-nodes are created for reference partitions, their assignments are predetermined and fixed throughout the partition optimization process^55^ using the `is_membership_fixed` argument of the `leidenalg.Optimiser().optimise_partition` method. As a result, query nodes that are similar to any of the existing reference partitions will be assigned to the corresponding group, while nodes further away will be assigned to new partitions. The resolution parameter γ will also control the level of alignment between query nodes and reference partitions: higher γ will lead to scattered query nodes, while lower γ will force the alignment.

### Predictive Modeling for Partition Extension

An alternative approach to extend existing partitions to unseen samples is through constructing a predictive model and applying it for inference. This is demonstrated in two experiments in Fig. 4A and **Supplementary Note 1**. In these experiments, we used manual annotations of reference samples to train gradient boosted tree classifiers, which were subsequently applied to unseen query samples.

The training dataset was constructed using the same set of reference nodes and used manual annotations as labels. The input contained reference node features, as well as 1-hop aggregated node features, which were computed by averaging features from the center nodes and their immediate spatial neighbors (defined by distance thresholding). This augmentation was inspired by the UTAG and SpaGCN methods and allowed the model to have larger fields of view. The test inputs for the query nodes were formulated similarly, and the trained models were employed to infer their cluster/partition assignments.

We evaluated a range of common machine learning methods including logistic regression, linear SVR, k-nearest neighbor classifier, and random forest. Gradient boosted trees implemented via XGBoost^56^ yielded the best performance.

### Evaluation Metrics

On the DKD Kidney dataset, DLPFC dataset, and Lung IMC dataset, we employed manual annotations to assess the performances of various unsupervised annotation tools. The following metrics were applied:

- Adjusted Rand Index^57^ (ARI) and Rand Index (RI): ARI and RI are measures that evaluate the similarity between two data clusterings, in which ARI also takes into account the probabilities of random agreement between two clusterings. ARI ranges from −1 to 1, where 1 indicates perfect agreement and 0 indicates a random agreement. We adapted the implementation in `scikit-learn`^58^: `sklearn.metrics.adjusted_rand_score.` RI ranges from 0 to 1, where 1 indicates perfect agreement. RI is only used in the Lung IMC dataset to reproduce metrics reported in its source publication. We adapted the implementation in scikit-learn: sklearn.metrics.rand_score.
- Homogeneity Score: homogeneity score measures if all points within one unsupervised cluster are members of a single label class. It ranges from 0 to 1, where 1 indicates perfect agreement. We adapted the implementation in `scikit-learn`: `sklearn.metrics.homogeneity_score.`
- F1 Score: F1 is an accuracy measure calculated as the harmonic mean of precision and recall. For each label compartment, the F1 score is calculated through the following process:

○ Labels: 1s are assigned to all nodes affiliated with the target label compartment, 0s are assigned to the rest.
○ Predictions: For a given partition, predictions are calculated using the indicator function of whether a node is assigned to that specific partition. A series of predictions will be derived for all partitions identified by the unsupervised annotation tool.
○ Metrics: Multiple F1s are calculated based on the labels and the series of predictions. The highest F1 score corresponds to the partition that has the best match with the target label compartment, and this score is taken as the final score.

## Acknowledgements

J.Z. is supported by NSF CAREER 1942926 and a Chan-Zuckerberg Biohub Investigator Award.

## Data and Code Availability

Datasets used in this manuscript are available at:

- DLPFC: https://research.libd.org/spatialLIBD/
- Lung IMC: https://doi.org/10.5281/zenodo.6376766
- DKD Kidney, UCSF Derm, TR Kidney: datasets are available for download and visualization at https://app.enablemedicine.com/portal/atlas-library

SCGP was implemented using emObject^59^ (https://docs.enablemedicine.com/emobject/) and Annotated data^60^ (https://anndata.readthedocs.io/en/latest/). The source code is publicly available at the following URL: https://gitlab.com/enable-medicine-public/scgp

## Supplementary Notes

### 1. Identify Tissue Structures in DLPFC with Partial Ground Truth

In the DLPFC four-sample joint partitioning experiment, among all the methods that generated unsupervised partitions/clusters, SpiceMix and SCGP-Extension achieved top performances with ARI over 0.6 (**Supplementary Fig. 2A**). To further improve annotation accuracy, we attempted utilizing the reference-query extension pipeline as well as the predictive modeling strategy with partial ground truth inputs.

This experiment simulated the application scenario in which we first acquire manual annotations on one specific sample from a dataset and then generalize it across all the remaining samples. One approach is to train a predictive model using the ground truth annotations and apply it to the rest. We adapted the gradient boosted tree method implemented in the XGBoost package and trained it with annotations from a representative sample (151673). Prediction results on other samples were well-aligned (**Supplementary Fig. 2B**, XGB Prediction), with a joint ARI of 0.64, surpassing all the unsupervised methods.

We further applied the reference-query extension pipeline using partial ground truth annotations, which follows the exact same steps as SCGP-Extension but using ground truth annotations instead of SCGP partitions as references. Two variants were tested, with different layers as references (**Supplementary Fig. 2B**, Label-Extension columns): Label_1_-Extension used all layers, while Label_2_-Extension excluded Layer 4. Results demonstrated better performances than SCGP-Extension and XGB Prediction (**Supplementary Fig. 2C**), achieving a joint ARI of 0.69.

Notably, characterization of the two thinner layers (Layer 2 and Layer 4, downward triangles and squares in **Supplementary Fig. 2D**) were worse in most methods, including ones that utilize partial ground truth. Layer 4 was especially unstable across samples, suggested by the noisy predictions on test samples from the XGB model. In fact, excluding Layer 4 in the reference yielded better ARI performances for Label_2_-Extension.

### 2. Annotate Transplant Rejection Kidney Samples with SCGP

We conducted CODEX imaging on a series of kidney tissue samples obtained from patients who experienced rejection responses after kidney transplantation^61^ (TR Kidney dataset, **Methods**). These tissues exhibit heavily deformed native kidney structures and substantial inflammation and immune cell infiltration. To assess the broader applicability of SCGP and SCGP-Extension, we performed the following analysis to identify and annotate common tissue structures across samples in this cohort.

We ran SCGP on a subset of ten samples with heavy rejection responses, encompassing approximately 363k cells (**Supplementary Fig. 3A**). The selection of these samples was based on considerations of data quality and computational efficiency. The primary SCGP run yielded 6 major partitions corresponding to kidney tissue structures including tubules, glomeruli, and blood vessels. Remarkably, despite the TR Kidney dataset being imaged using a different biomarker panel than the DKD Kidney dataset, we successfully derived the same set of tissue structures using different protein biomarker signatures (**Supplementary Fig. 3C**). Particularly in the case of glomeruli, the absence of partition-specific biomarkers (e.g., CCR6, Nestin) did not impair SCGP’s ability to recognize these structures. Theoretically, the differences in the expression patterns of multiple proteins are reflected by the vector distances between cells, which enable SCGP to isolate the glomeruli from their surrounding tissues. An additional partition (red) exhibiting high expression of immune cell biomarkers including CD45, CD20, and CD4 suggested the presence of substantial amounts of immune cells.

Subsequently, we attempted extending the primary partitions to a group of control samples with minimal immune responses, comprising five samples and around 110k cells. These samples were not well-integrated using SCGP (**Supplementary Fig. 6B**), where the same tissue structures from different samples were assigned to separate partitions. On the contrary, SCGP-Extension consistently recognized the same set of tissue structures (Fig. 4D and **Supplementary Fig. 3B**). We further noticed major differences between the two conditions: normal samples displayed much denser spatial organization of tubules and higher biomarker expression (e.g., CD31, CD34) in glomeruli.

Notably, we observed unsuccessful partitions in one sample of the TR Kidney dataset, potentially due to the uneven background signals of several biomarkers (**Supplementary Fig. 6A-C**, bottom row). Unsupervised annotation tools tend to highlight the most prominent differences between nodes, which, in this specific sample, come from the background artifact (gray partition in **Supplementary Fig. 3B**). SCGP-Extension, instead, enforces external references of known tissue structures, thereby downweighting the influences of unwanted signals. As a result, we successfully identified consistent tissue structures using SCGP-Extension (**Supplementary Fig. 6D**).

### 3. Annotate Healthy Lung Samples with SCGP

We employed SCGP and various unsupervised annotation tools to analyze a cohort of healthy lung specimens imaged using IMC (Lung IMC dataset, **Methods**). The dataset was adopted from a recent study^15,50^ and consisted of 26 samples with 28 biomarkers. Samples were manually annotated for major anatomical structures including airway, alveolar space, connective tissue, vessels, submucosal glands, and cartilage.

We first applied all unsupervised annotation tools to the combination of all 26 samples in a joint partitioning manner. Among all tested methods, UTAG, SpiceMix, and SCGP aligned better with manual annotations (**Supplementary Fig. 4A**). We further evaluated the alignment scores for each sample (Fig. 4F), and SCGP exhibited top performance with a median ARI of 0.464, followed by SpiceMix (median ARI=0.416) and UTAG (median ARI=0.364). Two additional clustering metrics presented by Kim et al.^15^: homogeneity score and rand index were also calculated and visualized (**Supplementary Fig. 4B-C**), showing similar outcomes.

However, even though partitions in each individual sample aligned well with manual annotations, the same tissue structures from different samples were often disconnected, as indicated by the different coloring of partitions in **Supplementary Fig. 4A**. To address this issue, we applied SCGP to a well-integrated subset comprising seven samples, in which tissue structures were consistently recognized across samples. Using this subset as references, we extended their partitions to the remaining samples. Notably, two manually annotated anatomical structures: submucosal glands and cartilage were absent in the reference samples. During extension, we successfully isolated them as unseen partitions not attached to any of the existing tissue structures (red and purple nodes in Fig. 4E, **Supplementary Fig. 4D**). As a result, SCGP-Extension demonstrated the best agreement with manual annotations and achieved significantly higher ARI scores (median ARI=0.564, *P* < 0.001, Wilcoxon signed-rank test).

### 4. Annotate Skin Samples of Different Disease Conditions with SCGP

We further assessed SCGP using a collection of four experiments on skin samples with different disease conditions (UCSF Derm dataset, **Methods**). These samples were imaged using CODEX in four separate experiments employing different biomarker panels. We extracted the shared biomarkers and performed unsupervised annotations using SCGP.

We conducted the initial SCGP experiment on 17 samples comprising 365k cells from experiment 1 (**Supplementary Table 1**). Major tissue structures including the epidermis layers, dermis layer, immune aggregates, and glands were recognized (**Supplementary Fig. 5A, E**). Next, we attempted both joint partitioning and reference-query extension on the remaining samples from other experiments. Both UTAG and SCGP experienced difficulties in integrating samples when applied in a joint partitioning manner, assigning different partitions to epidermis layers from different samples. This was likely due to the systematic differences between experiments (**Supplementary Fig. 7A-B**). SCGP-Extension, instead, delivered satisfactory results, recognizing consistent tissue structures across experiments. (**Supplementary Fig. 5B-D and Supplementary Fig. 7D**).

It is visually evident that samples of different disease conditions exhibit distinct spatial organizations of tissue structures. We aimed to further investigate the differences and correlate different disease conditions with tissue structure-level interpretations. To achieve this, we proposed heuristic metrics based on tissue structure annotations and assessed them across samples. We first derived the lengths and thicknesses of the epidermis layers using the corresponding annotations. Based on the coordinates of all the nodes assigned to the epidermis partition, we created contours surrounding these nodes. Lengths and thicknesses of epidermis layers were calculated treating the contours as masks (Fig. 5D). By comparing epidermis thicknesses across samples, we noticed that atopic dermatitis samples had significant thickenings (*P* < 0.001, two-sided two-sample t-test) of epidermis, showing a wide distribution of thicknesses (Fig. 5F). In the other metric, we quantified the density of immune aggregates by deriving the area proportion of the immune aggregate partitions. Results suggested significant enrichment of immune cells in atopic dermatitis samples compared to normal samples^47^ (*P* < 0.00, two-sided two-sample t-test).

### 5. Annotate mIF Images as Spatial Grids of Patches with SCGP

For spatial proteomics measurements (CODEX and IMC), we primarily discussed applications of SCGP to spatial graphs of cells in the main text, which require the preprocessing step of cell segmentation. In various scenarios where cell segmentation is unavailable or fails to meet expectations, an alternative approach would be defining spatial units that are not reliant on cells. We here present an example application of SCGP to samples from the DKD Kidney dataset, employing small square patches as spatial units for the graph community detection.

To begin, we dissected the mIF images into small square patches through a sliding window mechanism (**Supplementary Fig. 8A**), which transformed the images into spatial grids of image patches. We then naturally established graph representations of the grids by treating each patch as a node, with its feature encapsulating the summarized protein biomarker expression. Spatial edges were constructed between immediately adjacent patches, and feature edges were created between patches that shared similar expression profiles. This treatment shares similarities with a Visium spatial transcriptomics dataset, as it examines the tissue sample using a regular grid. The resolution of the grid can be adjusted by varying the size of the patches. In this particular experiment, the side length of each patch was set to 12 µm, similar to the diameter of a cell.

We conducted joint partitioning using SCGP on three representative samples of different DKD classes (**Supplementary Fig. 8B-C**), results aligned remarkably well with the manual annotations of healthy and mild DKD samples, identifying the same set of tissue structures. In the severe DKD (IIB) sample, the fibrotic glomeruli were not recognized as blood vessels (as in cell-based SCGP) or as a separate partition (as in SCGP-Extension) shown in Fig. 4A. Instead, they were assigned as combinations of the basement membrane and normal glomeruli partitions, suggesting the degradation of normal glomerular structures into scar tissues. Additionally, patch-based SCGP recognized a distinct immune aggregates partition that is enriched in CD45, CD68, and DAPI (i.e., higher cell density) expression (**Supplementary Fig. 8C**). This partition was also spatially more abundant in the severe DKD sample, agreeing with our understanding of DKD progression.

### 6. Running Time of Unsupervised Annotation Methods

We estimated running time for all unsupervised annotation methods on two major tasks:

- Joint clustering/partitioning of 17 samples from the DKD Kidney dataset, containing 137,654 cells;
- Joint clustering/partitioning of 4 samples from the same specimen (Br8100) from the DLPFC Visium dataset, containing 14,364 spots.

The time profiling is performed on an amazon cloud service ec2 instance (r6i.16xlarge or g4dn.16xlarge if GPU is required). See **Supplementary Table 2** for full results.

Among all the tested methods, CN, UTAG, SCGP, and SCGP-Extension had the shortest running time, comparable to common clustering algorithms. SpaGCN had slightly longer running time owing to its network optimization procedure; it still completed the DKD Kidney experiment within 10 minutes. Spatial LDA, BayesSpace and SpiceMix required substantially more time to finish the optimizations, posing considerable challenges in hyperparameter tuning.

For unsupervised annotation methods benchmarked in this work, we searched the hyperparameter space (e.g., resolution parameter, number of clusters) for each method and chose the best-performing results that produced 6-7 clusters. Below we briefly described the pipelines for these methods, please refer to their source publications and code bases for full details.

- KMeans

○ Input: Biomarker expression vectors of cells/spots;
○ Clustering: KMeans algorithm, implemented by scikit-learn^58^.
- Leiden^38^

○ Input: *k*-nearest neighbor (*k* = 15) graph constructed using the biomarker expression vectors of cells/spots;
○ Clustering: Leiden algorithm, implemented by leidenalg (https://github.com/vtraag/leidenalg).
- Cellular Neighborhood (CN)^13,26^

○ Input: Cell types are first identified through leiden clustering. For each cell, a composition (frequency of cell types) vector is calculated based on a window of 20 nearest neighboring cells (including the center cell) as measured by Euclidean distance between X/Y coordinates.
○ Clustering: KMeans algorithm, implemented by scikit-learn.
○ Not applied to the DLPFC dataset due to requirements of cell types.
- UTAG^15^

○ Input: For each cell/spot, an average biomarker expression vector is calculated over all neighboring cells/spots and the center cell/spot within a window surrounding the center cell/spot thresholded by Euclidean distance (18μm in DKD Kidney dataset), referred to as the spatially aggregated feature matrix. *k* -nearest neighbor (*k* = 15) graph is then constructed using the aggregated expression vectors.
○ Clustering: Leiden algorithm, implemented by leidenalg.
- Spatial LDA^27^

○ Input: Cell types are first identified through leiden clustering. For each cell, its local environment is encoded as the count of cell types (bag-of-cell) within a window surrounding the center cell thresholded by Euclidean distance (20μm in DKD Kidney dataset). Spatial prior (i.e., adjacency between cells) is first constructed by computing the Voronoi partitioning of cell positions, in which pairs of cells that share a facet in the Voronoi partitioning are connected, then reduced to a minimum spanning tree based on the edges.
○ Clustering: Latent Dirichlet Allocation with spatial prior, implemented in https://github.com/calico/spatial_lda.
○ Not applied to the DLPFC dataset due to requirements of cell types.
- BayesSpace^28^

○ Input: Top principal components of the log transformed and normalized gene expression counts.
○ Clustering: Spots are modeled using a fully Bayesian model with a Markov random field prior, specified by the Potts model. Model parameters are estimated using a Markov chain Monte Carlo method. We adapted codes (in R) from https://edward130603.github.io/BayesSpace/articles/BayesSpace.html.
○ Only used on the DLPFC data.
- SpaGCN^31^

○ Input: Biomarker expression vectors of cells/spots are reduced to their top 20 principal components and then constructed into a weighted undirected graph, in which edges are weighted by Euclidean distances (histology information is not included in this study).
○ Clustering: A one-layer graph convolutional network on the input graph generates initial embeddings for cells/spots, which are clustered using the Louvain algorithm. Network parameters and cluster centroids are optimized by minimizing a soft assignment-based loss function using stochastic gradient descent with momentum until convergence. We adapted codes from https://github.com/jianhuupenn/SpaGCN.
- SpiceMix^24^

○ Input: a Hidden Markov Random Field model is constructed based on the graphical model, which treats cells/spots as nodes and connects spatially adjacent pairs (through Delaunay triangulation) with edges.
○ Clustering: Cells/spots are first clustered using the Louvain algorithm to initialize estimates of hidden states and model parameters, which are further iteratively optimized via coordinate ascent. We adapted codes from .

## Supplementary Figures

**Supplementary Figure 1.**
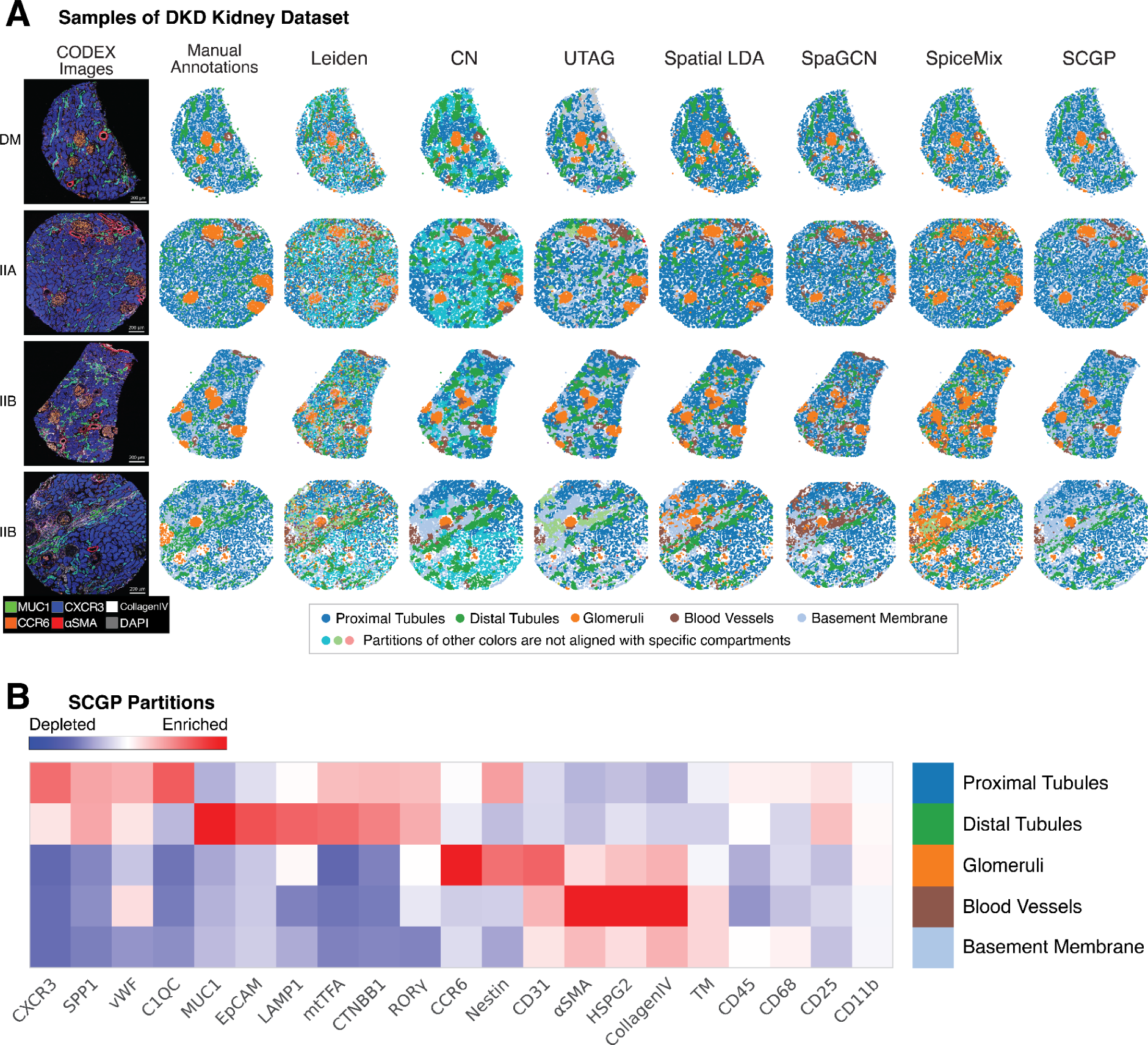
Additional examples and results on the DKD Kidney dataset. **A.** Clustering/partitioning outputs from unsupervised annotation tools on four samples of different DKD classes are illustrated. Qualitatively, Leiden clustering identified cell types that are not spatially smooth; CN defined an extra cluster (cyan) for proximal tubules; UTAG defined multiple smaller clusters (light green, gray, purple) that do not correspond to any compartments; Spatial LDA and SpiceMix had more noisy annotations for glomeruli (orange); SpaGCN misrecognized some regions as blood vessels; SCGP misrecognized some fibrotic glomeruli as blood vessels. **B.** Full heatmap for all biomarkers tested in the DKD Kidney dataset shows signature protein biomarkers for SCGP partitions, with each partition corresponding to a manually annotated compartment.

**Supplementary Figure 2.**
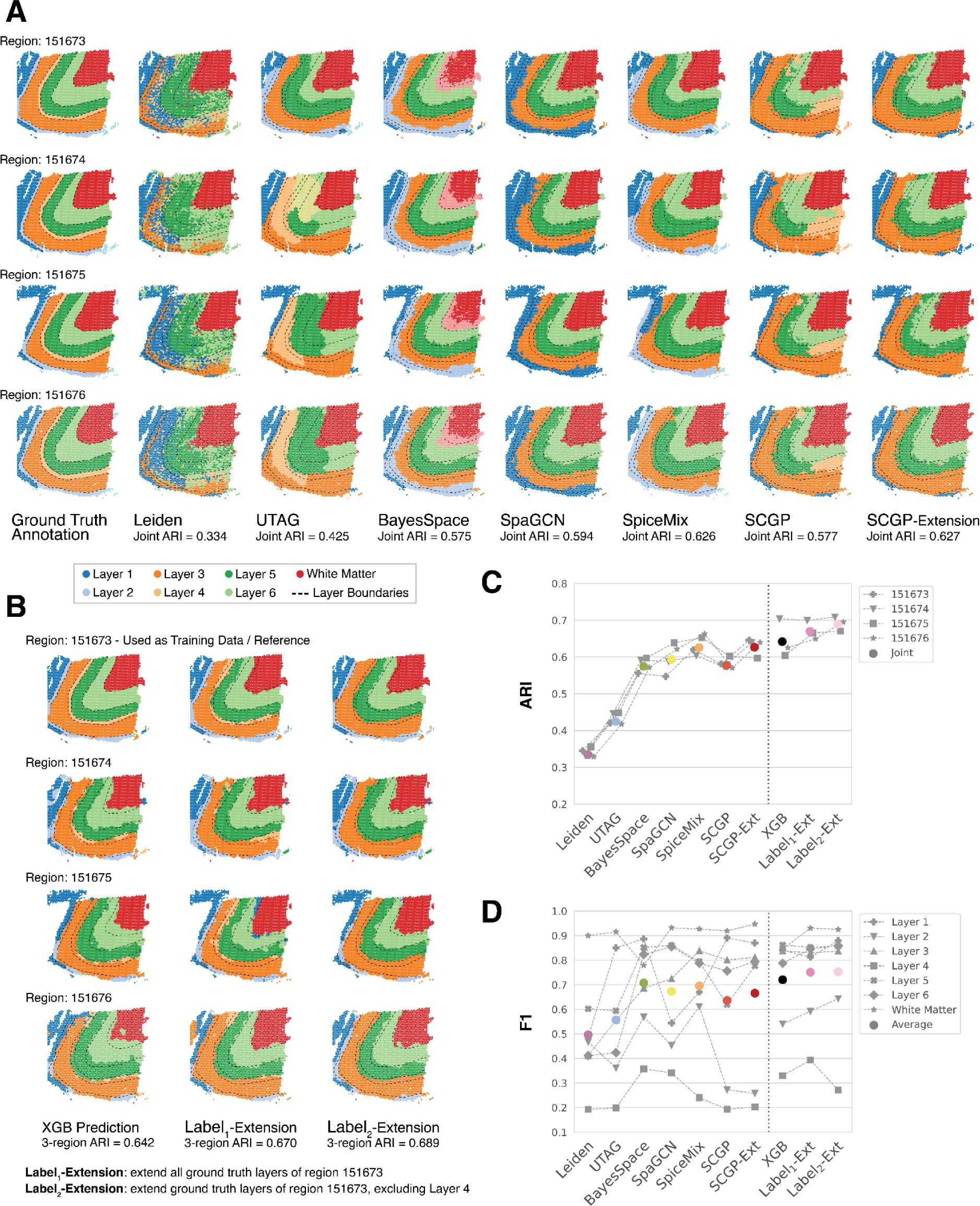
Joint partitioning of four DLPFC samples. **A.** Joint unsupervised annotations recognized major layers in four DLPFC samples. **B.** Supervised annotations and reference-query extensions utilizing labels of one sample (the top row) yielded better outcomes. Note that Layer 4 cannot be robustly recognized even with partial ground truth labels. **C.** ARIs are calculated between joint annotations and ground truth labels for each sample (gray markers) and the joint of all samples (colored circles). Methods utilizing partial ground truth yielded better performances, in which extension performs better than XGBoost predictions. **D.** F1s are calculated between joint annotations and ground truth labels on each of the 7 layers (gray markers), and average values over all layers are plotted as colored circles. Note the accuracy for Layer 2 and Layer 4 are constantly worse.

**Supplementary Figure 3.**
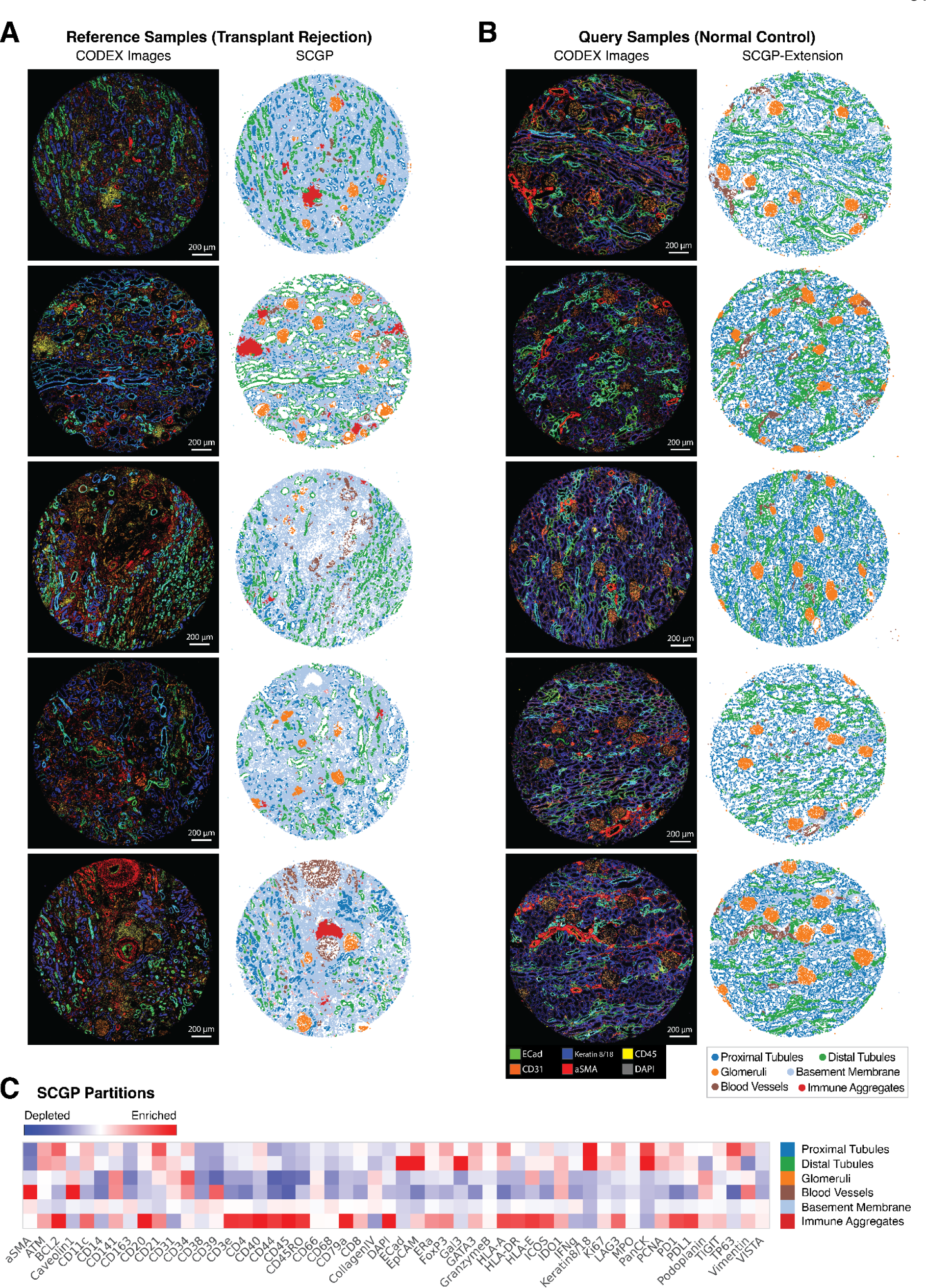
SCGP annotations of TR Kidney dataset. **A.** Primary SCGP experiment on samples with heavy inflammation defined six major partitions. **B.** Partitions were extended to normal samples that have minimal inflammation. Note the absence of the immune aggregates partition and the denser arrangement of tubules and glomeruli. **C.** Signature protein biomarker expression for the six tissue structures identified by SCGP are visualized in the heatmap.

**Supplementary Figure 4.**
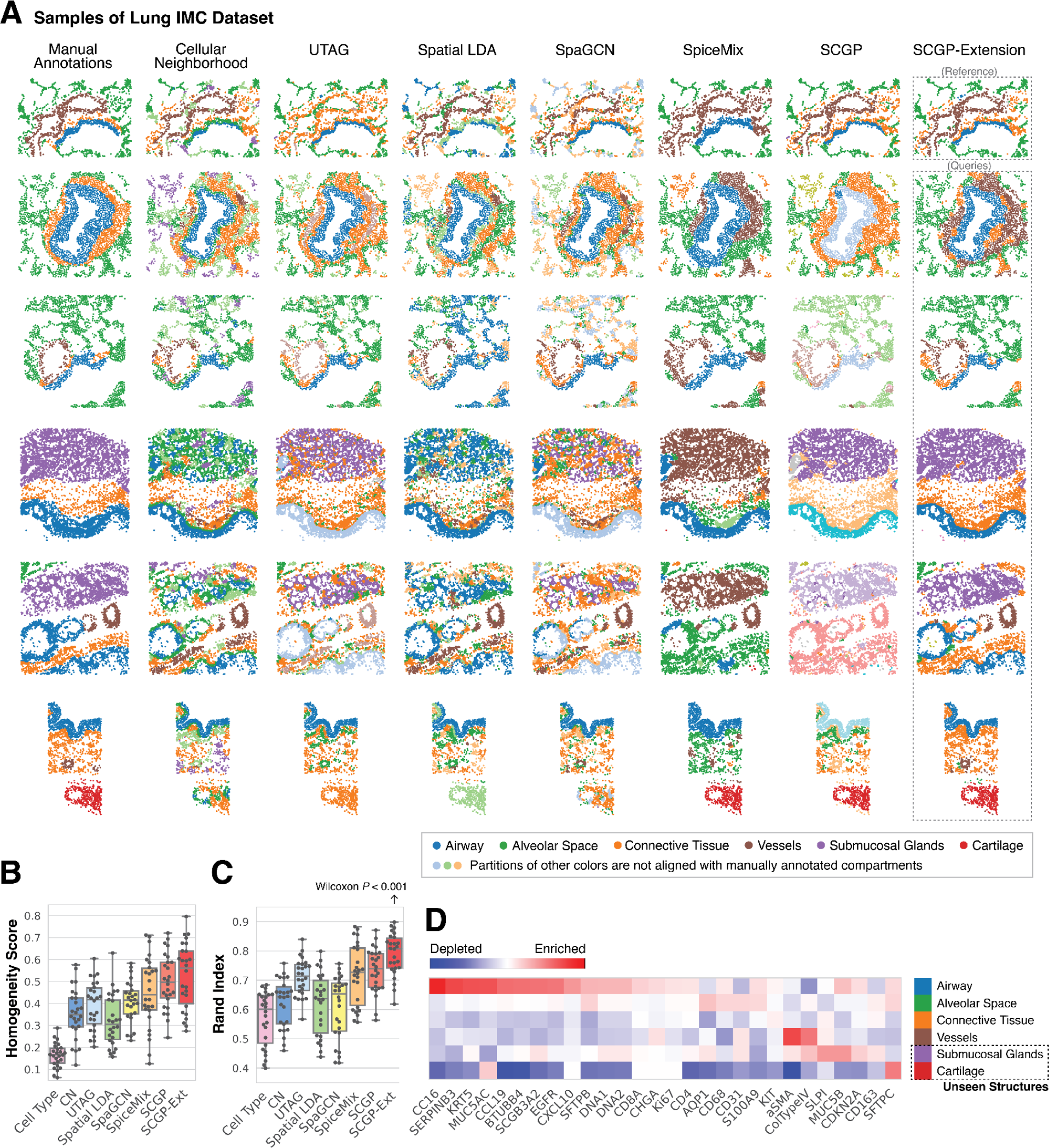
SCGP annotations of Lung IMC dataset. **A.** Clustering/partitioning outputs from various unsupervised annotation tools on representative lung samples are visualized. SCGP-Extension achieved the best visual alignment with manual annotations. **B.** and **C.** Metrics^15^ (homogeneity score and rand index) are calculated between unsupervised annotations and manual annotations. **D.** Signature protein biomarker expression for the tissue structures identified by SCGP-Extension are visualized in the heatmap, corresponding to the six manually annotated compartments. Note that submucosal glands and cartilage are two tissue structures only identified in the query samples.

**Supplementary Figure 5.**
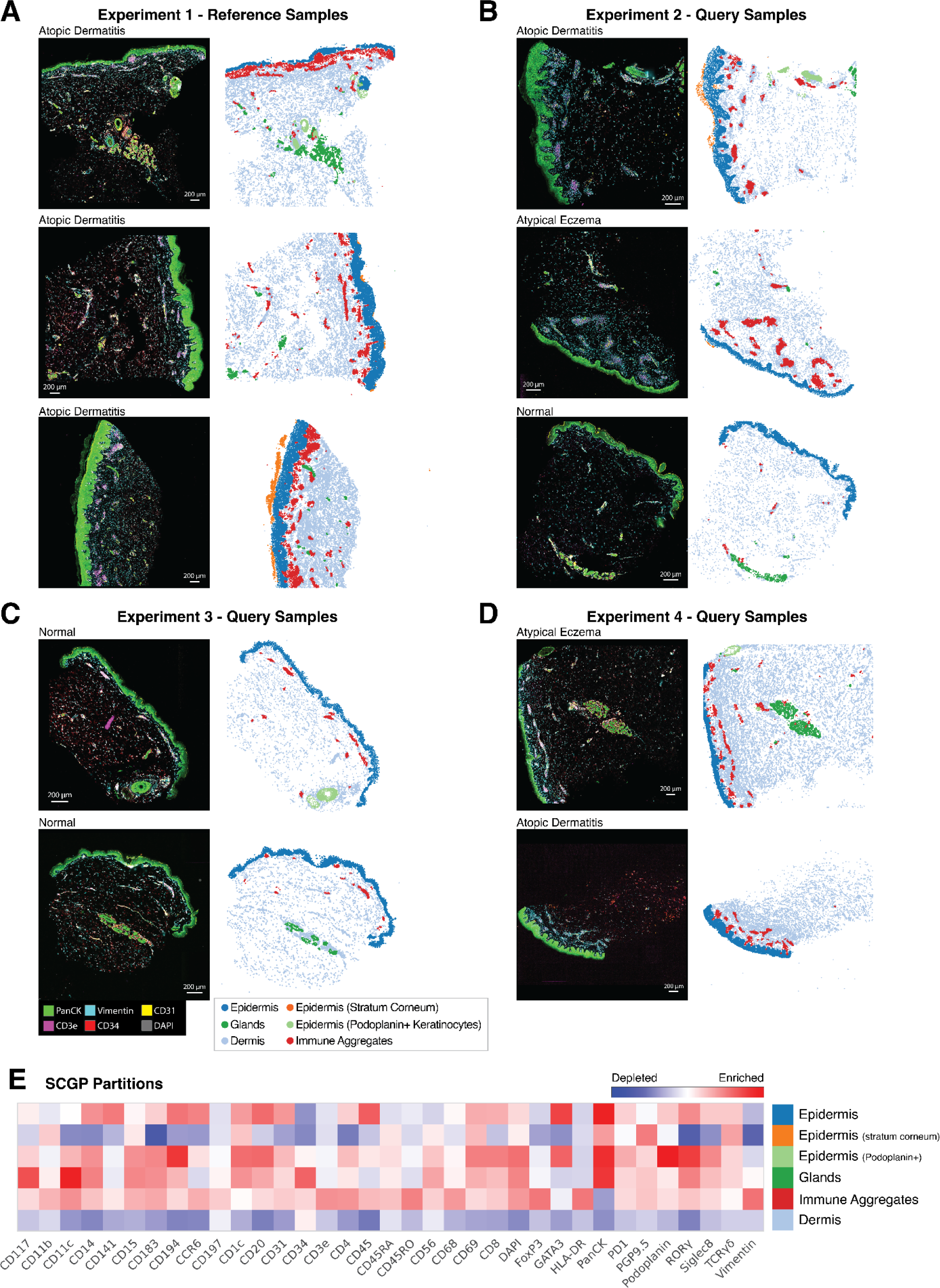
SCGP annotations of UCSF Derm dataset. **A.** Primary SCGP annotations on samples from experiment 1 defined major partitions. **B**-**D.** Partitions were extended to samples from different experiments that have different skin conditions **E.** Signature protein biomarker expression for the major partitions identified by SCGP are visualized in the heatmap.

**Supplementary Figure 6.**
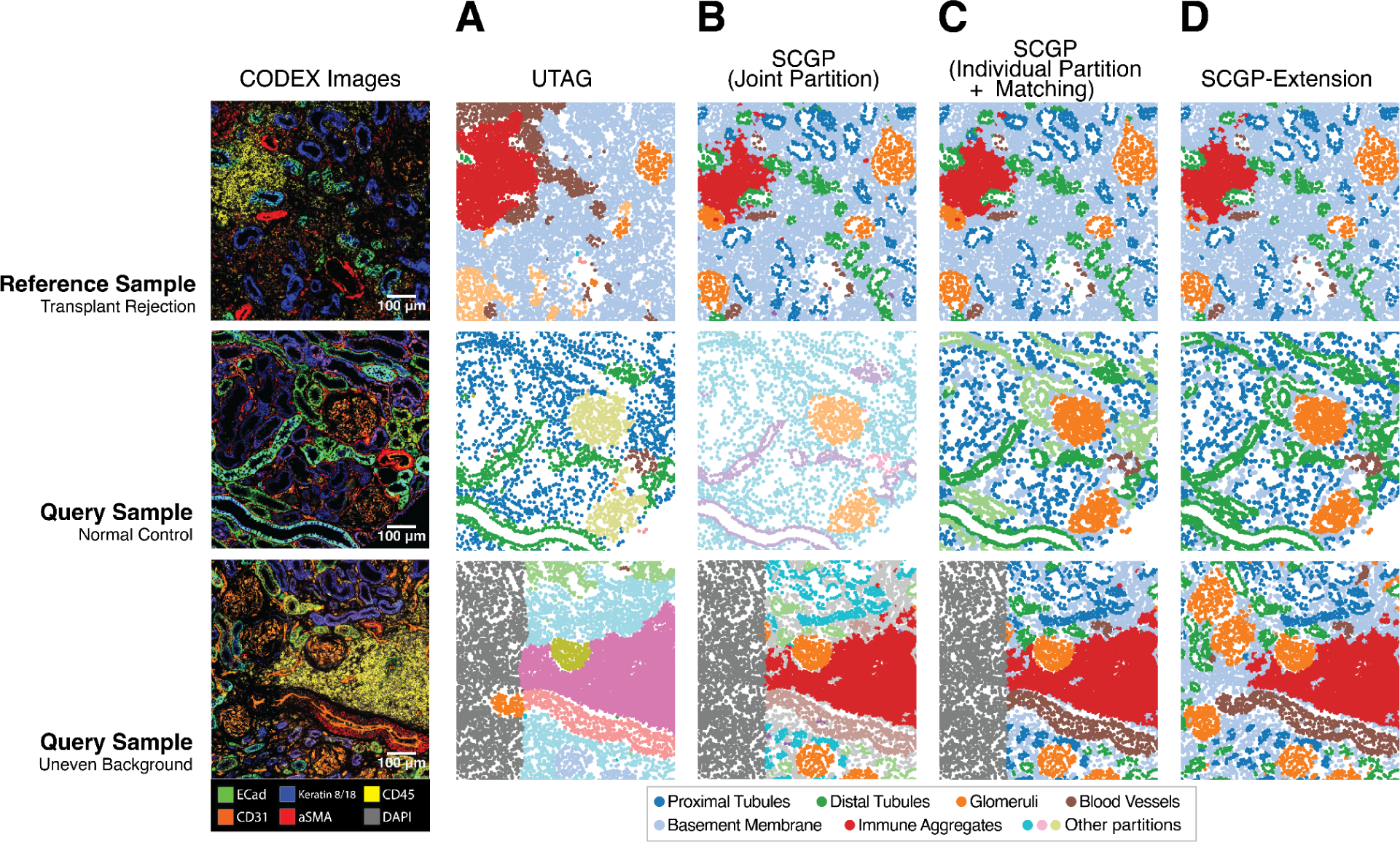
Annotating representative samples from TR Kidney dataset using variants of SCGP. **A.** UTAG defined partitions do not correspond well to tissue structures. **B.** SCGP failed to assign structures of the same type into the same partitions. Note the different coloring of samples. **C.** Individual SCGP partitions of different samples were matched post hoc to reflect shared tissue structures. Note that the uneven background interfered with partitions. **D.** SCGP-Extension enabled consistent recognition of tissue structures regardless of artifact (uneven background) and disease condition differences.

**Supplementary Figure 7.**
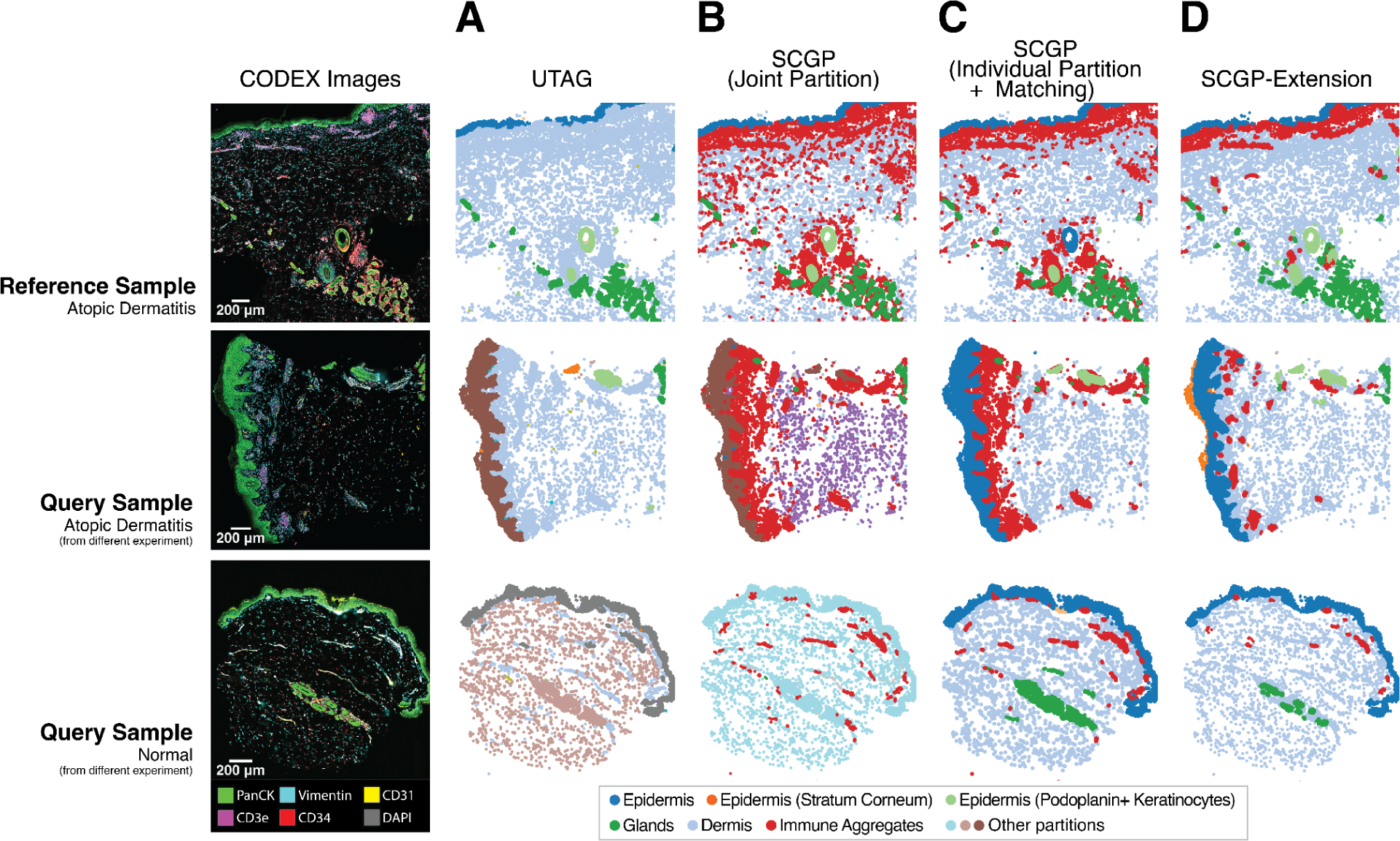
Annotating representative samples from UCSF Derm dataset using variants of SCGP. **A.** and **B.** Joint partitioning with UTAG and SCGP failed to assign structures of the same type into the same partitions. Note the different coloring of epidermis in the samples. **C** Individual partitions of different samples are matched post hoc to reflect shared tissue structures. **D.** Reference-query extension pipeline enabled consistent recognition of tissue structures across samples from multiple experiments with different skin conditions.

**Supplementary Figure 8.**
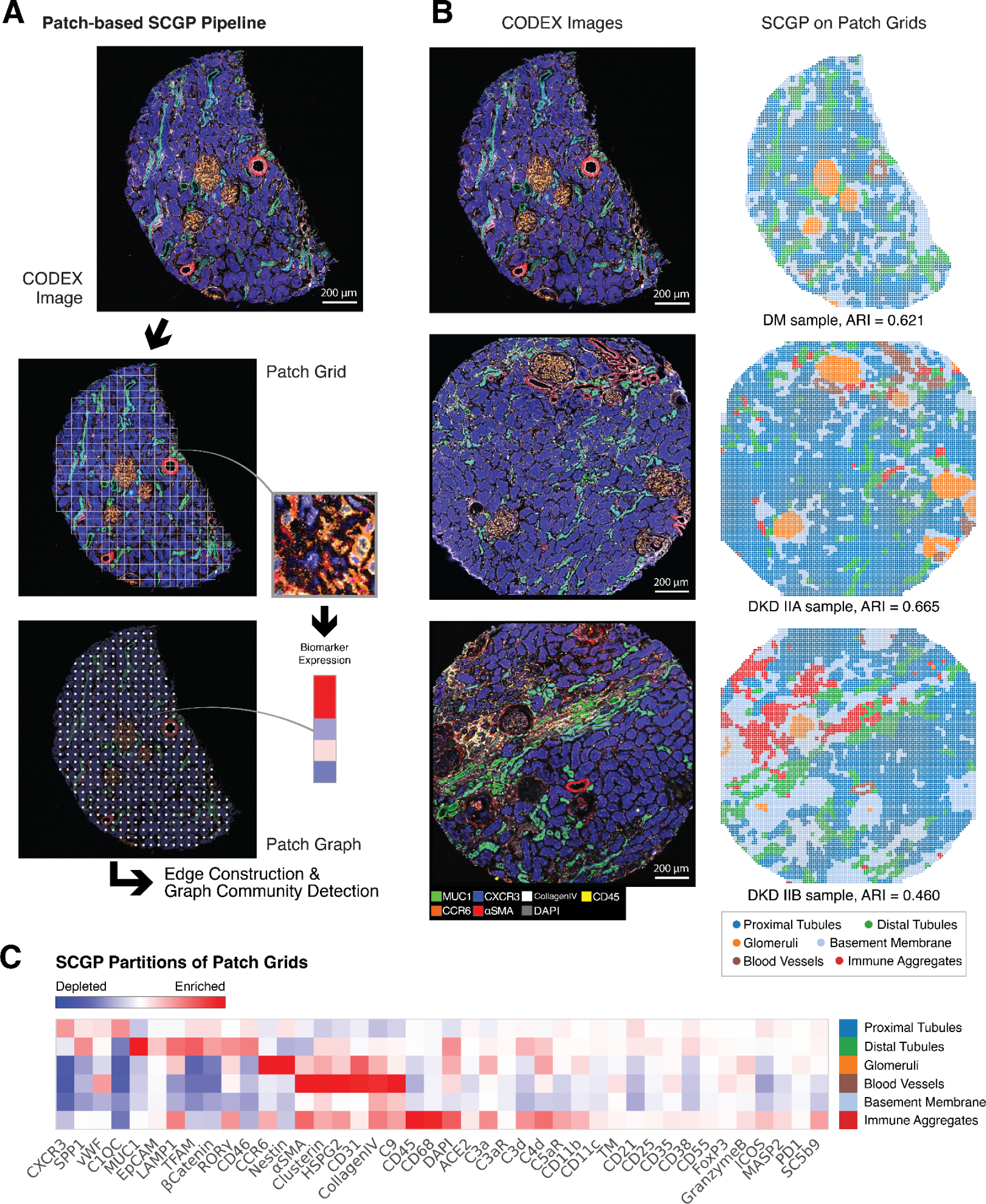
SCGP annotations of mIF images as spatial grids of patches **A.** mIF image was first dissected into small patches through a sliding window. Each patch was summarized as a feature vector representing biomarker expression within the field of view. Spatial graph of patches was constructed as input to SCGP. **B.** SCGP on the patch grids of three mIF images yielded similar results as cell-based SCGP, recognizing tubules, glomeruli, blood vessels, and immune aggregates in the tissue. **C.** Signature protein biomarker expressions are visualized in the heatmap for the partitions identified by patch-based SCGP.

## Supplementary Tables

**Supplementary Table 1.** Details of datasets: number of samples/cells/patients and major grouping or patient characteristics.

| Dataset | N (samples) | N (cells/spots) | N (patients) | Major groups / patient characteristics |
| --- | --- | --- | --- | --- |
| DKD Kidney | 17 | 137,654 | 12 | Kidney sections with different DKD classes:<br>• Healthy kidneys from diabetic individuals (DM): 7 sections<br>• DKD class IIA: 2 sections<br>• DKD class IIA-B: 3 sections<br>• DKD class IIB: 4 sections<br>• DKD class III: 1 section |
| DLPFC | 12 | 47,681 | 3 | Postmortem DLPFC samples from three independent neurotypical adult donors |
| TR Kidney | 22 | 765,129 | 22 | Kidney samples sorted into two groups:<br>• Case group: 17 samples (transplant rejection)<br>• Control group: 5 samples (normal) |
| Lung IMC | 26 | 69,830 | 3 | Lung samples from three healthy donor lung specimens: sections of airways extending from proximal bronchi and succeeding divisions to terminal and respiratory bronchioles. |
| UCSF Derm | 44 | 588,867 | 33 | Skin samples with different skin conditions:<br>• Atopic Dermatitis (AD)<br>• Normal (N)<br>The dataset is collected from 4 experiments:<br>• Experiment 1: 22 samples (22 AD)<br>• Experiment 2: 3 samples (3 N)<br>• Experiment 3: 11 samples (4 AD, 4 N)<br>• Experiment 4: 8 samples (2 AD, 3 N) |

**Supplementary Table 2.**
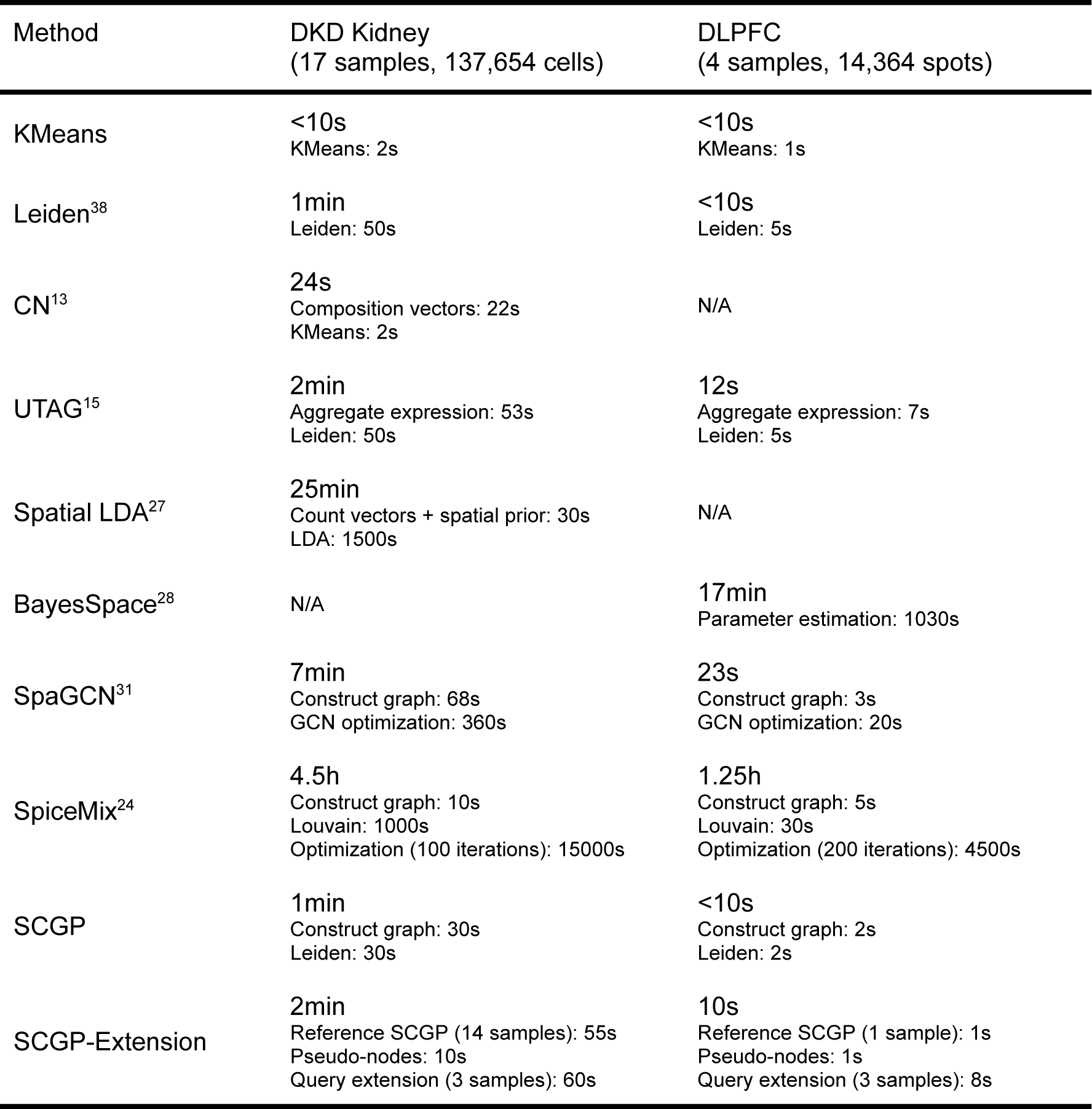
Running time of unsupervised annotation methods.

| Method | DKD Kidney<br>(17 samples, 137,654 cells) | DLPFC<br>(4 samples, 14,364 spots) |
| --- | --- | --- |
| KMeans | <10s<br>KMeans: 2s | <10s<br>KMeans: 1s |
| Leiden<sup>38</sup> | 1min<br>Leiden: 50s | <10s<br>Leiden: 5s |
| CN<sup>13</sup> | 24s<br>Composition vectors: 22s<br>KMeans: 2s | N/A |
| UTAG<sup>15</sup> | 2min<br>Aggregate expression: 53s<br>Leiden: 50s | 12s<br>Aggregate expression: 7s<br>Leiden: 5s |
| Spatial LDA<sup>27</sup> | 25min<br>Count vectors + spatial prior: 30s<br>LDA: 1500s | N/A |
| BayesSpace<sup>28</sup> | N/A | 17min<br>Parameter estimation: 1030s |
| SpaGCN<sup>31</sup> | 7min<br>Construct graph: 68s<br>GCN optimization: 360s | 23s<br>Construct graph: 3s<br>GCN optimization: 20s |
| SpiceMix<sup>24</sup> | 4.5h<br>Construct graph: 10s<br>Louvain: 1000s<br>Optimization (100 iterations): 15000s | 1.25h<br>Construct graph: 5s<br>Louvain: 30s<br>Optimization (200 iterations): 4500s |
| SCGP | 1min<br>Construct graph: 30s<br>Leiden: 30s | <10s<br>Construct graph: 2s<br>Leiden: 2s |
| SCGP-Extension | 2min<br>Reference SCGP (14 samples): 55s<br>Pseudo-nodes: 10s<br>Query extension (3 samples): 60s | 10s<br>Reference SCGP (1 sample): 1s<br>Pseudo-nodes: 1s<br>Query extension (3 samples): 8s |

